# Osh6 requires Ist2 for localization to the ER-PM contacts and efficient phosphatidylserine transport

**DOI:** 10.1101/2020.01.31.928440

**Authors:** Juan Martín D’Ambrosio, Véronique Albanèse, Nicolas-Frédéric Lipp, Lucile Fleuriot, Delphine Debayle, Guillaume Drin, Alenka Čopič

## Abstract

Osh6 and Osh7 are lipid transfer proteins (LTPs) that move phosphatidylserine (PS) from the endoplasmic reticulum (ER) to the plasma membrane (PM). High PS level at the PM is key for many cellular functions. Intriguingly, Osh6/7 localize to ER-PM contact sites, although they lack membrane-targeting motifs, in contrast to multidomain LTPs that both bridge membranes and convey lipids. We show that Osh6 localization to contact sites depends on its interaction with the cytosolic tail of the ER-PM tether Ist2, a homologue of TMEM16 proteins. We identify a motif in the Ist2 tail, conserved in yeasts, as the Osh6-binding region, and we map an Ist2-binding surface on Osh6. Mutations in the Ist2 tail phenocopy *osh6Δ osh7Δ* deletion: they decrease cellular PS levels, and block PS transport to the PM. Our study unveils an unexpected partnership between a TMEM16-like protein and a soluble LTP, which together mediate lipid transport at contact sites.

## Introduction

Many lipids are non-uniformly distributed between the membranes of eukaryotic cells, thus conferring to organelle membranes their distinct properties and identities (Harayama and Riezman, 2018). Lipid transfer proteins (LTPs) contribute to this uneven distribution by transporting specific lipid species between compartments. A prominent eukaryotic LTP family are the oxysterol-binding protein–related proteins (ORP) and the related Osh proteins, which are characterized by a conserved lipid-binding domain (ORD, for oxysterol-binding protein– related domain) (Raychaudhuri and Prinz, 2010). Crystal structures revealed that ORDs could accommodate one lipid molecule within their binding pocket, either sterol or phosphatidylserine (PS) (Im et al., 2005; Maeda et al., 2013). Importantly, many ORDs have been shown to alternately encapsulate phosphatidylinositol-phosphates (PI4P or PI(4,5)P_2_), and sequence analyses suggest that PI4P is the common ligand for all ORP/Osh proteins. This dual lipid specificity allows ORP/Osh proteins to operate lipid exchange between compartments and use phosphoinositide metabolism to transport the second lipid species against its concentration gradient (Chung et al., 2015; de Saint-Jean et al., 2011; Filseck et al., 2015; Ghai et al., 2017; Mesmin et al., 2013; Moser Von Filseck et al., 2015).

We have been studying Osh6, which, together with its close paralog Osh7, transports PS from the endoplasmic reticulum (ER) to the plasma membrane (PM) in yeast (Maeda et al., 2013). PS is synthesized at the ER, but its concentration is much higher at the PM, specifically in the cytosolic leaflet, where it mediates recruitment of proteins involved in signaling and establishment of cell polarity, and is important for initiation of endocytosis (Kay and Fairn, 2019). We have shown that Osh6 exchanges PS with PI4P, which is synthesized at the PM and then hydrolyzed at the ER by the PI4P-phosphatase Sac1, and that PI4P transport and degradation are required for efficient PS transport in yeast (Moser Von Filseck et al., 2015). This is also true for the closest Osh6 homologues in mammalian cells, ORP5 and ORP8 (Chung et al., 2015).

In agreement with their proposed function, Osh6 and Osh7 can be observed at the cortical ER, which represents sites of close apposition between the ER and the PM (Schulz et al., 2009). These ER-PM contact sites are maintained by tethering proteins that are able to simultaneously bind to both compartments. A number of such proteins have been shown to maintain the ER-PM contacts in yeast (Manford et al., 2012). LTPs themselves often contain additional tethering domains or motifs that mediate their localization to contact sites (Wong et al., 2017). For example, ORP5 and ORP8 contain a transmembrane (TM) region that is embedded in the ER, and a pleckstrin homology (PH) domain that binds the PM (Olkkonen and Li, 2013). Many other LTPs, including several ORP/Osh proteins, bind to the ER via ER-resident VAP proteins using a short FFAT (two phenylalanines in an acidic tract) motif upstream of their ORD domain (Loewen et al., 2003; Murphy and Levine, 2016). In contrast, Osh6 consists only of an ORD and how it targets the ER and the PM is not known. We have previously demonstrated that to some extent, the cellular localization of Osh6 is regulated by its intrinsic avidity for lipid membranes (Lipp et al., 2019). However, this does not explain how Osh6 specifically targets its donor and acceptor compartments and how it maintains accuracy in PS transport.

Here, we show that the localization of Osh6 to cortical ER depends on its binding to the ER-PM tether Ist2. Ist2 is an intriguing tether that contains a long and disorder cytosolic tail and a TM region embedded in the ER, which shares homology with the TMEM16 proteins, a family of Ca^2+^-activated lipid scramblases (Brunner et al., 2014; Kralt et al., 2014; Manford et al., 2012; Wolf et al., 2012). We show that a short segment of the disordered tail is required for Osh6 binding and localization, and that this interaction is necessary for Osh6-mediated PS transport to the PM. Finally, we identify residues in a conserved region of Osh6, distal from the entrance to the lipid binding pocket, that are involved in binding to Ist2. Our results unveil an unexpected partnership between a TMEM16-like protein and a cytosolic LTP, which enables lipid flux at contact sites, and is important for PS homeostasis in yeast.

## Results

### Ist2 is required for Osh6 localization to the cortical ER

Many LTPs localize to the ER using a short FFAT motif that binds to the VAP proteins (Murphy and Levine, 2016). No such motif can be identified in the ORD of Osh6. We therefore speculated that another protein was responsible for Osh6 localization to the ER-PM contacts. To identify such protein(s), we endogenously tagged Osh6 with a TAP-tag and performed affinity chromatography followed by proteomic analysis using mass spectrometry (Fig. S1). Among the proteins that co-purified with Osh6 (Table S1), we identified Ist2, an important yeast ER-PM tether (Collado et al., 2019; Hoffmann et al., 2019; Manford et al., 2012; Wolf et al., 2012). Due to their over-lapping localization, Ist2 seemed a good candidate for mediating Osh6 targeting to the cortical ER. An interaction between these two proteins was also detected, but not further confirmed, in a high-throughput proteomics study (Babu et al., 2012). In agreement with these results, we found that deletion of *IST2* rendered Osh6 completely cytosolic (Fig. 1A). In contrast, Osh6 remained cortical in *scs2Δ scs22Δ* cells, which lack the two yeast VAP proteins that recruit multi-domain Osh proteins to contact sites, namely Osh1, Osh2 and Osh3 (Weber-Boyvat et al., 2014). Furthermore, when we tagged Osh6 and Ist2 at their carboxy termini with Venus N-ter (VN) and C-ter (VC) halves to perform bimolecular fluorescence complementation (BiFC), we could observe fluorescent signal at the cell cortex (Fig. 1B). This was not the case when we expressed Osh6-VN or Ist2-VC alone, or when we co-expressed Osh6-VN with a Ist2 truncated at aa 590, fused to VC. Importantly, the ratio between cortical and soluble Osh6 depended on the level of expression of Ist2 (Fig. 1C,D). We conclude that Ist2 is the limiting factor for Osh6 localization to ER-PM contacts and that the two proteins may directly interact.

**Fig. 1.**
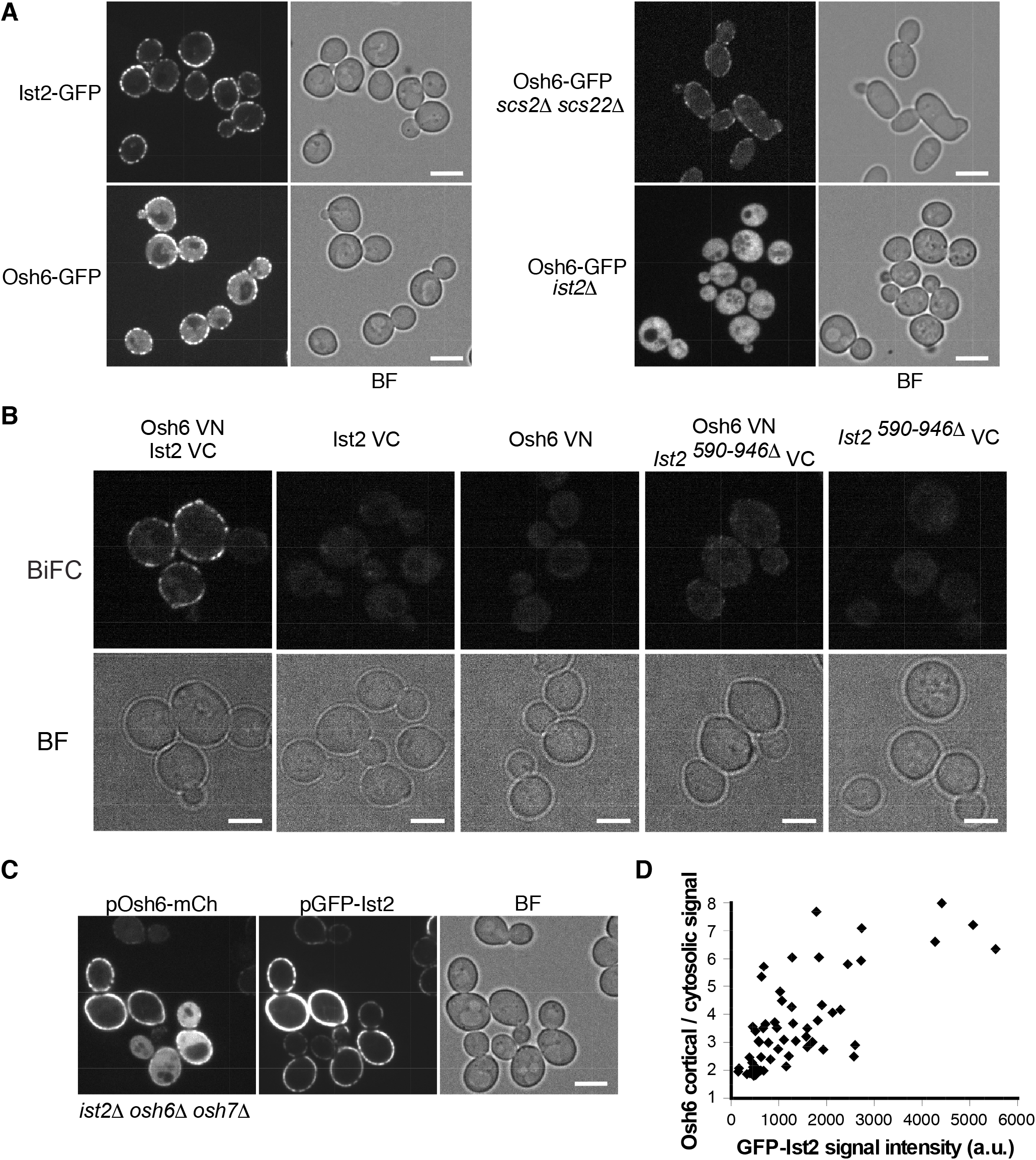
Ist2 is required for Osh6 localization to the cortical ER. **(A)** Localization of chromosomally-tagged Ist2-GFP and Osh6-GFP in WT, *scs2Δ scs22Δ*, or *ist2Δ* cells. **(B)** Bimolecular fluorescence complementation (BiFC) in diploid cells expressing endogenously tagged Osh6-VN and Ist2-VC or Ist2^590Δ^-VC (Ist2 truncated at aa590), as indicated. **(C)** Localization of Osh6-mCherry and GFP-Ist2 expressed from low-copy plasmids in *ist2Δ osh6Δ osh7Δ* cells. **(D)** The ratio of Osh6-mCherry peripheral vs. cytosolic signal in (C) was quantified as a function of GFP-Ist2 fluorescence (a.u.= arbitrary units). Each symbol represents one cell from the same experiment; two independent experiments were quantified, yielding similar results. All strains were imaged at least 3 times. Scale Bar = 5 μm. Bright field (BF).

### Osh6 interacts with the disordered cytosolic tail of Ist2

Ist2 is an intriguing ER-PM tether. It is embedded in the ER membrane via a domain containing 8 predicted TM helices, which bears structural homology to a large mammalian family of Ca^2+^-activated lipid scramblases and/or ion channels, called TMEM16 proteins (Brunner et al., 2014) (Fig. 2A). This domain contains a C-ter appendage of about 300 aa that is specific to fungi and is predicted to be largely disordered, followed by a polybasic motif that binds to the PM (Kralt et al., 2014; Maass et al., 2009). Interestingly, it was shown that the cytosolic tail of Ist2 could be significantly truncated (to more than half of its original length) without an effect on Ist2 localization (Kralt et al., 2014). In contrast, we observed that a longer part of the tail (from aa 696 to C-ter) was required for cortical localization of Osh6, in accordance with our BiFC data, suggesting that Osh6 may bind to a region in this portion (Fig. 1B and Fig. 2B). We tested this hypothesis using a yeast two-hybrid assay, in which we expressed full-length Osh6 as prey and different fragments of Ist2 as bait (Fig. 2C and Fig. S2A). We observed a strong interaction between Osh6 and C-ter, but not N-ter cytosolic fragment of Ist2, as well as with shorter fragments of the C-ter tail, down to 19 aa (aa 729-747). Similar results were obtained with Osh7, whereas Osh4/Kes1, another soluble Osh protein, did not show any interaction with Ist2 (Fig. 2C and Fig. S2B). We could also detect interaction between Osh7 and full-length Ist2 using the BiFC assay (Fig. S2C). We then mutated select residues within the Ist2[729-747] region, specifically the ones that may be phosphorylated. Substitution of T736 and T743 with alanine was sufficient to block interaction with Osh6 and to render it cytosolic (Fig. 2C,D), whereas mutation of other residues had a smaller or no effect on Osh6 binding (Fig. S2D). Alignment of TMEM16 homologous sequences from budding yeasts (Saccharomycotina) shows that the [729-747] region is highly conserved in the monophyletic clade containing Saccharomycodacae and Saccharomycetacea (Shen et al., 2018) (Fig. 3). Especially well conserved are the residues T736 and T743; in contrast, the sequence of the rest of the tail is not conserved.

**Fig. 2.**
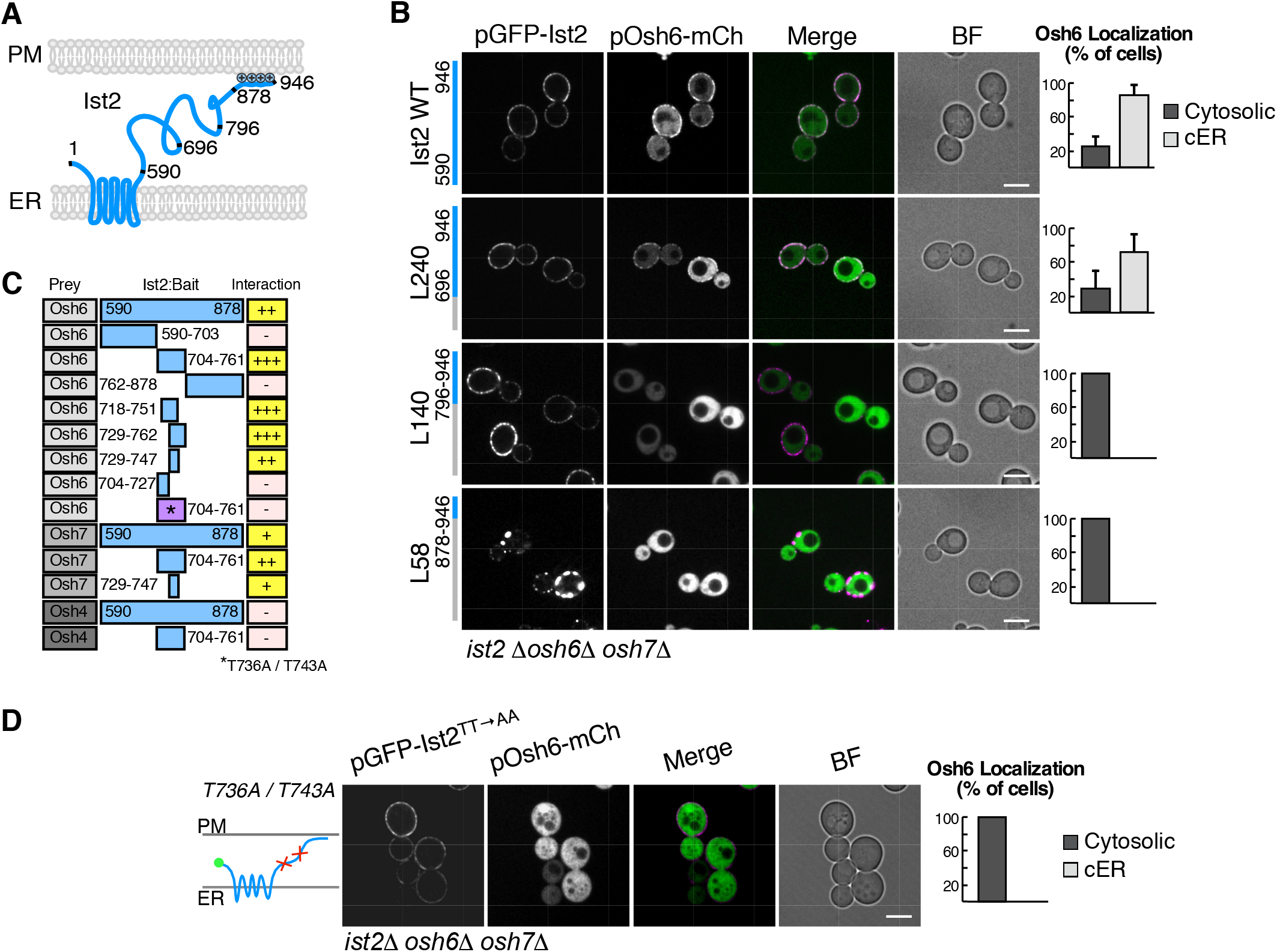
Osh6 interacts with the disordered cytosolic tail of Ist2. **(A)** Schematic representation of Ist2 predicted topology. Positions of aa in the disordered cytosolic tail are indicated, as well as the C-term polybasic domain (+). **(B)** Localization of Osh6-mCherry and GFP-Ist2 or deletion mutants (L240=Ist2^*590-696Δ*^, L140=Ist2^*590-796Δ*^ and L58=Ist2^*590-878Δ*^), expressed from plasmids in *ist2Δ osh6Δ osh7Δ* cells. Graphs show fraction of cells with visible cortical ER (cER) Osh6 signal vs. cytosolic only. The bars show mean+s.e.m. from 3 independent experiments (n=60 cells for each experiment). **(C)** Mapping of Osh6 interaction site on Ist2 by yeast two-hybrid assay. Osh6, Osh7 and Osh4 were used as prey and full length Ist2 cytosolic tail (aa 590-878) and shorter fragments, as indicated, were used as bait; relative interaction was scored from growth on reporter plates in 3 independent assays. (*) indicates T736A and T743A mutations. **(D)** Localization of GFP-Ist2^*T736A/T743A*^ and Osh6-mCherry, expressed from plasmids in *ist2Δ osh6Δ osh7Δ*. Quantification as in (B) (n=60 cells for each of 3 independent experiments). Scale Bar = 5 μm.

**Fig. 3.**
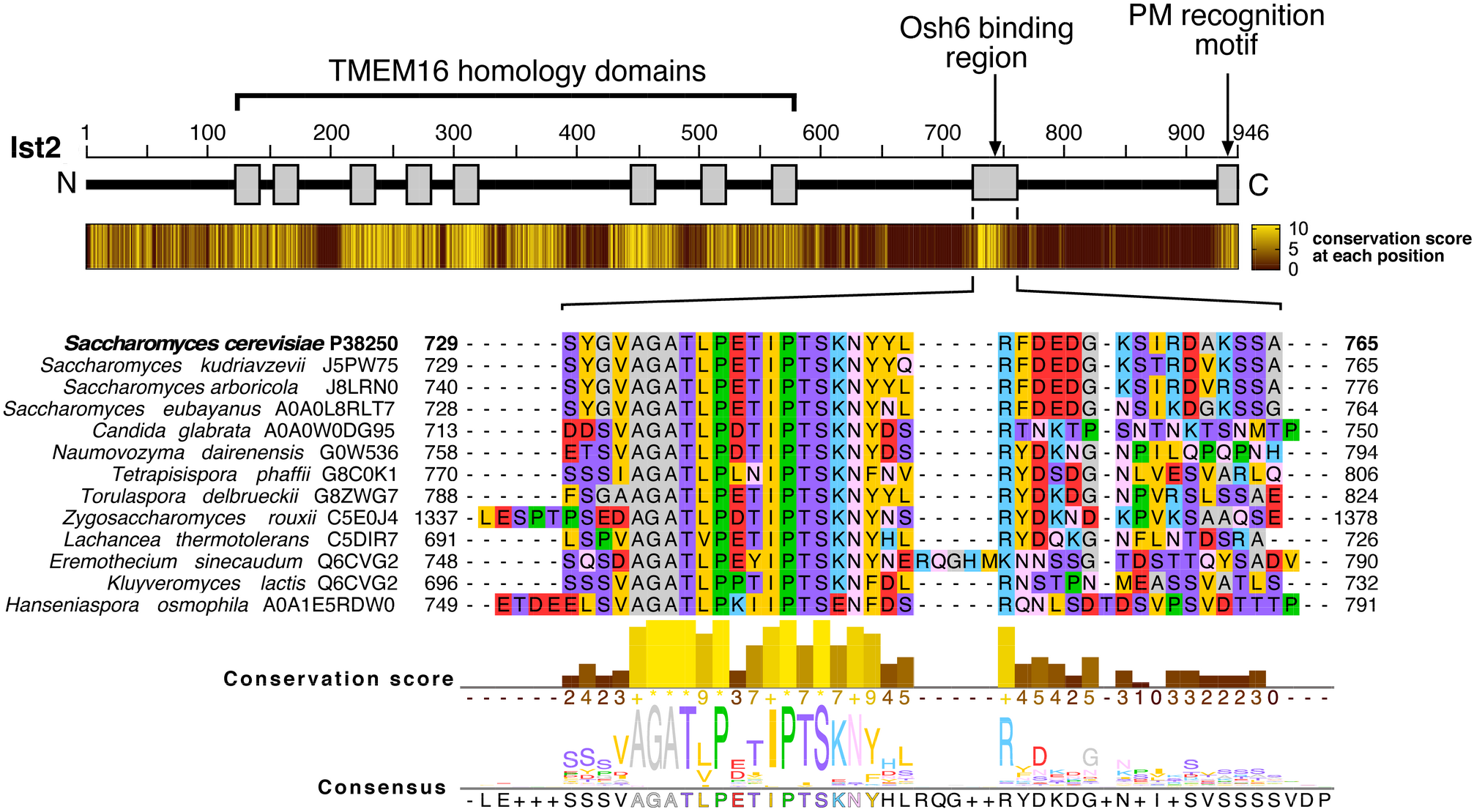
Detection of a conserved motif in the intrinsically disordered tether of TMEM16 orthologues in budding yeasts. Sequences of TMEM16 orthologues from 58 Saccharomycotina species were aligned. Conservation score from a subset of 34 sequences, which correspond to a monophyletic clade that includes *Saccharomycodacae* and *Saccharomycetacea* (Shen et al., 2018), is shown on a heatmap as a brown to yellow scale (0 to 11). Thirteen sequences of the conserved Osh6-binding region from the subset of species are shown, as well as the consensus Osh6-binding sequence from the whole subset. See Fig. S7 for the full alignment.

We next asked whether this conserved Osh6-interacting fragment of the Ist2 tail was sufficient for localizing Osh6 to the cortical ER. For this, we used a previously described Ist2 mutant Ist2-RL, in which the whole disordered tail sequence was randomized (Kralt et al., 2014). When co-expressed with Osh6 as fluorescent fusions in *ist2Δ* yeast, Ist2-RL still localized to the cortical ER, whereas Osh6 did not. However, insertion of Ist2[729-747] fragment into Ist2-RL partially rescued Osh6 cortical localization (Fig. 4A). Rescue was even more efficient with a slightly longer fragment (aa 718-751), mirroring the strong interaction detected in the yeast two-hybrid assay (Fig. 2C and Fig. 4B). Together, these results suggest that Osh6 localizes to ER-PM contact sites by binding to a short region within the disordered tail of Ist2. A more quantitative analysis where we compared the ratio of cortical to cytosolic Osh6 signal in a cell as a function of total Osh6 signal showed only a small difference between BFP-Ist2-RL^718-751WT^ and BFP-Ist2-expressing cells, suggesting that the rest of the tail at best has a small effect on Osh6 localization (Fig. 4C).

**Fig. 4.**
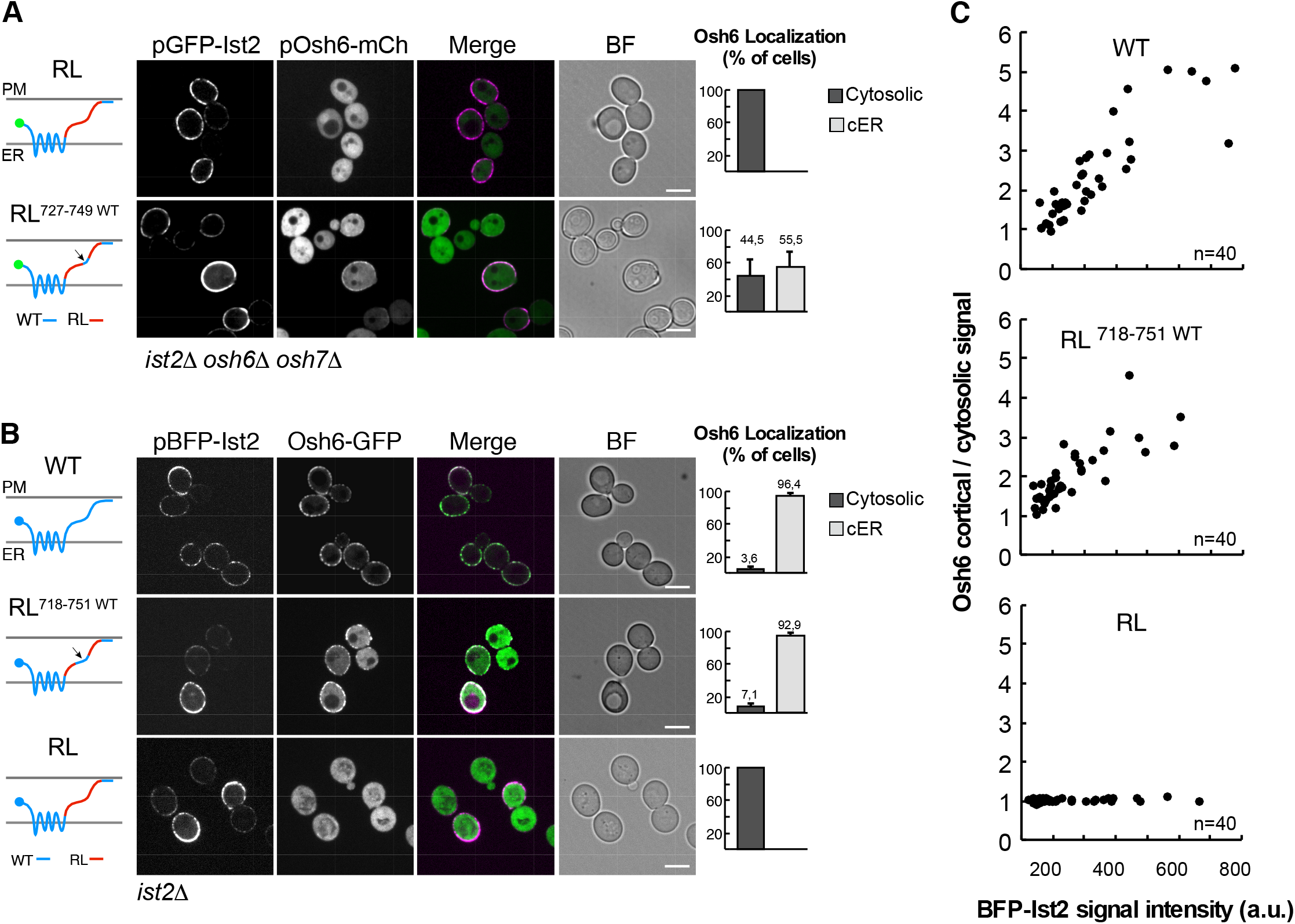
Osh6 binding site in Ist2 tail is sufficient for localizing Osh6 to the ER-PM contact sites. **(A**) Localization of GFP-Ist2-RL (randomized linker, aa 596-917) or GFP-RL^727-749WT^ (aa 727-749 in RL replaced with corresponding WT Ist2 sequence) and Osh6-mCherry, expressed from plasmids in *ist2Δ osh6Δ osh7Δ* cells. **(B)** Localization of plasmid-borne BFP-Ist2, BFP-Ist2-RL^718-751WT^ (aa 718-751 from Ist2 in RL) or BFP-Ist2-RL and chromosomally-tagged Osh6-GFP in *ist2Δ* cells. BFP = blue fluorescent protein. Graphs in A) and B) show fraction of cells with visible cortical ER (cER) Osh6 signal vs. only cytosolic signal. The bars show mean+s.e.m. from 3 independent experiments (n=60 cells for each experiment). **(C)** Quantification of Osh6 peripheral/cytosolic signal as a function of BFP-Ist2 fluorescence in BFP-Ist2 or BFP-Ist2-RL^718-751WT^ and BFP-Ist2-RL-expressing cells shown in (B). Each symbol represents one cell (n=40). Graph shows a representative of 3 independent experiments. Scale Bar = 5 μm.

### Ist2 is required for Osh6-mediated PS transport to the PM

Because Osh6 and Osh7 require Ist2 for their localization to ER-PM contacts, we tested whether this interaction was important for PS transport. We first evaluated the effect of *ist2* mutations on general lipid homeostasis in the cell. Lipidomic analysis of whole cell extracts showed that both *osh6Δ osh7Δ* and *ist2Δ* cells had a strong (about 2-fold) decrease in PS levels compared to WT, similar to the decrease in PS observed in a strain lacking all ER-PM tethers (Quon et al., 2018). In contrast, the levels of phosphatidylcholine (PC), phosphatidylethanolamine (PE) or phosphatidylinositol (PI) were not significantly affected (Fig. 5 and Fig. S3). Interestingly, two point mutations in the Ist2 tail that disrupt the Osh6/7 binding site (T736A/T743A substitution and Δ[736-743] deletion), but do not affect Ist2 localization or ER-PM tethering, led to a similar decrease in PS levels as *ist2Δ* (Fig. 2D and Fig. 5). These results suggest that the decrease in PS levels in *ist2* mutants, like in the *osh6Δ osh7Δ* mutant, is directly due to a block in PS export from the ER, and may result from a repression of PS synthesis due to elevated PS levels at the ER (Sohn et al., 2016; Tani and Kuge, 2014).

**Fig. 5.**
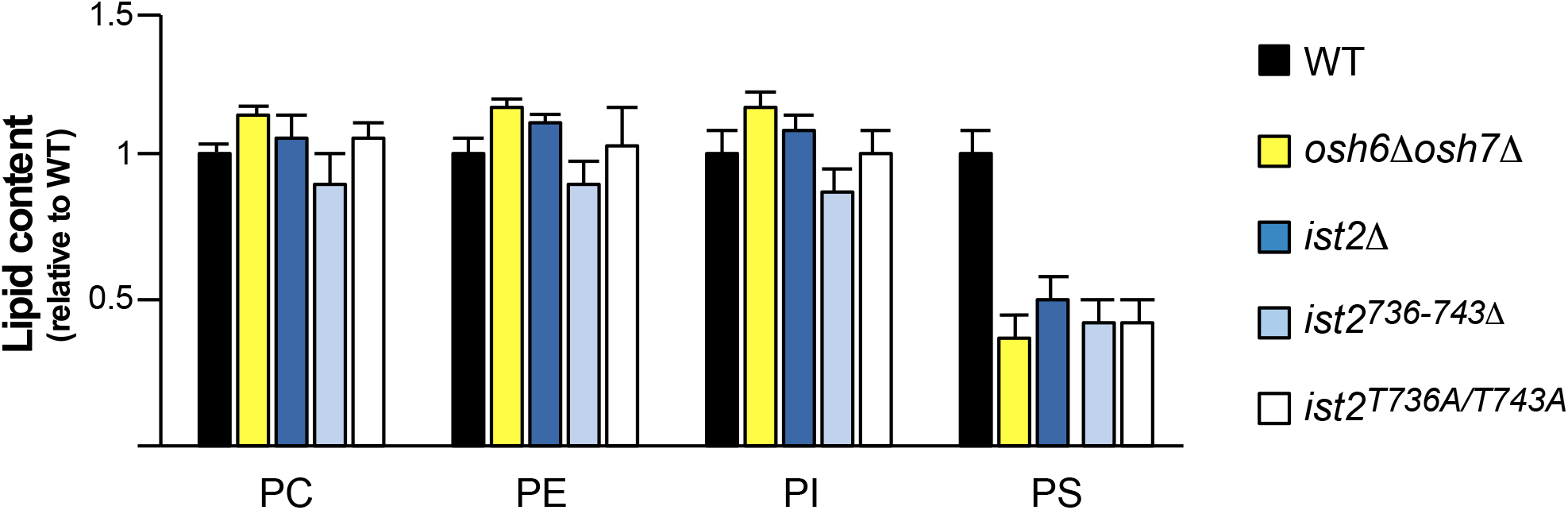
Mutations in Ist2 tail decrease steady-state levels of PS. Lipidomic analysis of WT, *osh6Δ osh7Δ, ist2Δ*, *ist2*^*736-743Δ*^ (chromosomal deletion of 8 codons) and *ist2*^*T736A/T743A*^ (chromosomal substitutions) cells. Lipid content is expressed relative to WT levels. Data are mean±s.d. from 3 independent samples. The experiment was repeated 2 or 3 times, depending on strain, with similar results.

To assess the influence of *ist2* mutation on PS transport from the ER, we analyzed the cellular PS distribution using the fluorescent PS reporter C2_Lact_-GFP (Fairn et al., 2011). We usually observed a decrease in the level of C2_Lact_-GFP at the PM in *osh6Δ osh7Δ* cells compared to WT, as previously reported (Maeda et al., 2013), as well as an effect of the deletion of *IST2.* However, there was variability between experiments and the difference was not always significant (Fig. S4). The absence of a strong effect on PS distribution at steady state is not surprising because it is known that the lack of Osh6 and Osh7 can be compensated by other, poorly understood mechanisms (Kay and Fairn, 2019; Ma et al., 2018).

To more precisely assess the involvement of Ist2-tail in PS transport, we used a cellular PS transport assay. For this, we used *cho1Δ* cells, which lack the PS synthase Cho1 and contain no PS, resulting in a cytosolic distribution of C2_Lact_-GFP (Fairn et al., 2011). Exogenous addition of lyso-PS, which is taken up by the cell (Riekhof et al., 2007; Spira et al., 2012), leads to PS synthesis via acylation at the ER and transport of PS to the PM over a time-course of 10-20 min, when Osh6/7 are functional (Maeda et al., 2013; Moser Von Filseck et al., 2015) (Fig. 6A,B). However, this was not the case in *cho1Δ ist2*^*736-743Δ*^ cells, in which PS accumulated at the ER, as demonstrated by the distribution of C2_Lact_-GFP, similar to *cho1Δ osh6Δ osh7Δ* cells. The small deletion in the Ist2 cytosolic tail therefore functionally phenocopied the absence of Osh6 and Osh7 for transport of newly-synthesized PS. We could fully rescue the PS transport defect in these cells by adding a plasmid-borne copy of *IST2* or of the chimeric construct *IST2-RL^718-751WT^*, which encodes Ist2 with randomized linker region except for the Osh6-interacting sequence aa718-751 (Fig. 6C,D). In contrast, expression of Ist2-RL or of a short soluble peptide Ist2[705-762], which contains the Osh6-interacting region, did not rescue the PS transport defect of *cho1Δ ist2*^*736-743Δ*^ cells. We conclude that localization of Osh6 to the ER-PM contact sites *via* its binding to the conserved region in the Ist2 tail is required for PS transport from the ER to the PM. This is likely also true for the closely related Osh7.

**Fig. 6.**
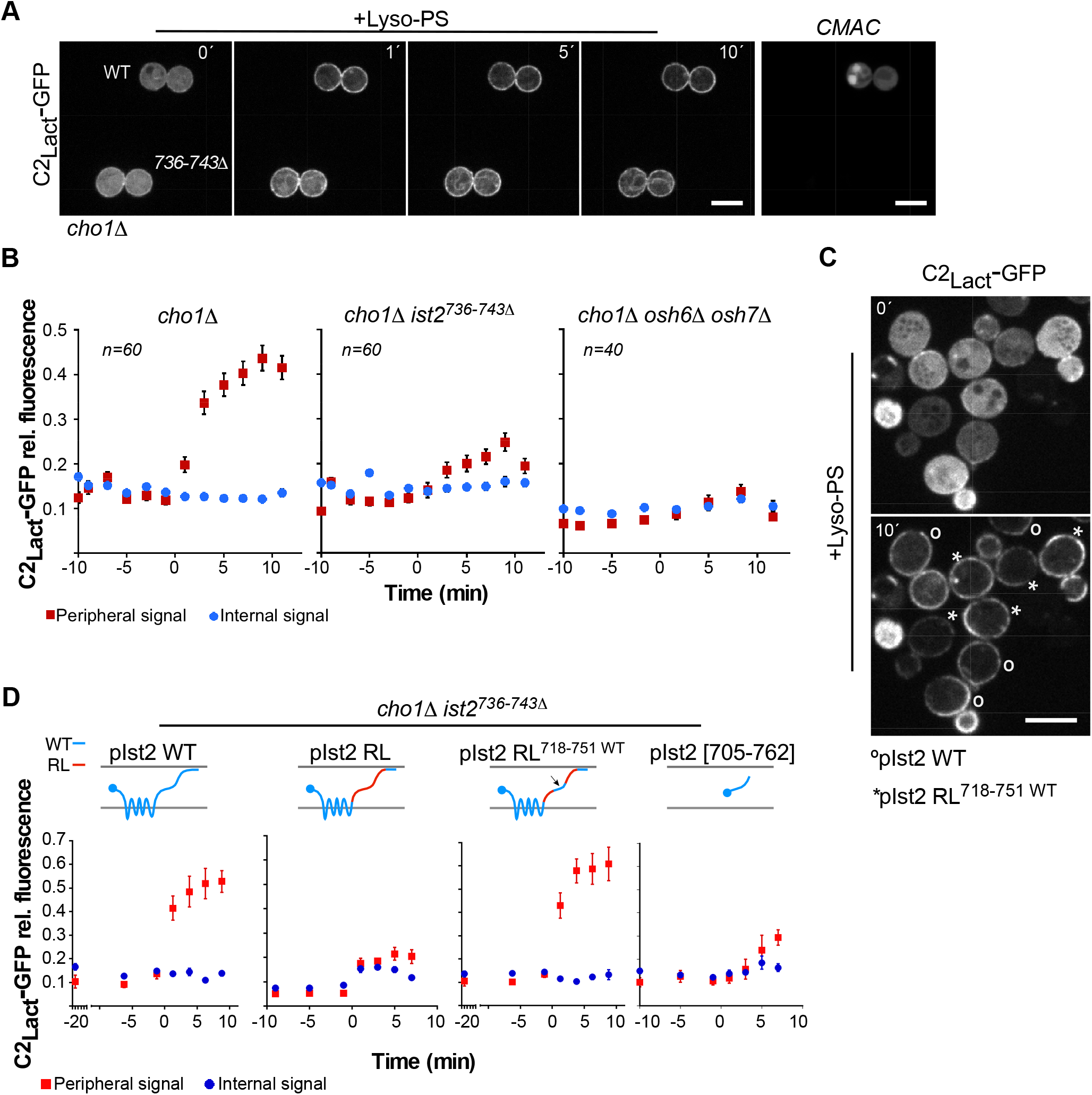
Binding of Osh6 to the disordered tail of Ist2 is required for PS transport to the PM. **(A)** Redistribution of C2_Lact_-GFP in *cho1Δ ist2736-743Δ* yeast, which lacks endogenous PS, after addition of 18:1 lyso-PS, compared to a *cho1Δ IST2-WT* control, imaged simultaneously. The control strain was labelled with the vacuolar dye CMAC (right panel) prior to imaging. Images were taken every 2 min. The first panel (t = 0’) shows the last time-point before C2_Lact_-GFP signal transition. The absolute timing of this event varied between experiments (10-20 min) due to instability of the lyso-PS suspension. **(B)** Quantification of C2_Lact_-GFP fluorescence over time from experiments like (A), with control and a mutant strain imaged simultaneously, in *cho1Δ, cho1Δ ist2*^*736-743Δ*^ and *cho1Δ osh6Δ osh7Δ*. Peripheral signal (red) = ER + PM; internal (blue) = mostly ER. Data are mean ± SEM from 3 independent experiments (20 cells per experiment). **(C)** C2_Lact_–GFP fluorescence in *cho1Δ ist2736-743Δ* cells with plasmid-borne BFP-Ist2 WT (°) or pBFP-Ist2 RL^718-751WT^ (*) after addition of lyso-PS at the indicated time points. (°) cells were labelled with CMAC (not shown). **(D)** Quantification C2_lact_-GFP fluorescence as in (B) in *cho1Δ ist2*^*736-743Δ*^ cells expressing Ist2 (WT), Ist2-RL, Ist2-RL^718-751WT^ or Ist2 fragment aa705-762. Data are mean±s.e.m. (n=15) from a representative of 3 independent experiments. Scale Bar = 5 μm.

### Mapping the Ist2 interaction site on Osh6

Our experiments suggest that Osh6 interacts with the cytosolic tail of Ist2, probably in a direct manner, although we cannot exclude the existence of an intermediate adaptor. We wanted to map the site on Osh6 that is responsible for the interaction with Ist2. Previous work revealed key residues (H157/H158 and L69) that coordinate the lipid ligand inside the binding pocket, and the importance of the N-ter region (1-69), which forms a lid over the binding pocket and regulates Osh6 interaction with lipid membranes (Lipp et al., 2019; Maeda et al., 2013; Moser Von Filseck et al., 2015). Mutation or deletion of these residues did not affect Osh6-Ist2 interaction (Fig. S5A,B). Using the yeast two-hybrid assay, we screened 29 new mutants of Osh6 with substitutions in surface-exposed and/or more highly conserved residues (Fig. 7A). Among these mutants, one with substitution of two adjacent aa located on the opposite side to the entry of the lipid-binding pocket, D141 and L142, to alanine showed no interaction with Ist2 and rendered Osh6-mCherry cytosolic (Fig. 7B and Fig. S5C). The same was true for Osh6(D141A/L142D), whereas mutation of only one residue (D141A, D141K or L142A), or a weaker substitution (D141K/L142I) were not sufficient to block the Ist2-Osh6 interaction (Fig. S5D). We analyzed the evolution of the Ist2 tail sequence and found that the Osh6-interacting region appears in a monophyletic subgroup of budding yeasts that includes Saccharomycodacea and Saccharomycetaceae (Fig. 7C). Interestingly, comparison with evolutionary conservation of Osh6 sequences reveals the apparition of a conserved region at the surface of the protein that includes D141 and L142 in this yeast subgroup (Fig. 7D). We found an even higher conservation in this region between Osh6 homologues in the whole genome duplication clade, after the Osh6/Osh7 duplication event, in line with our observation that Osh6 and Osh7 both interact with Ist2.

**Fig. 7.**
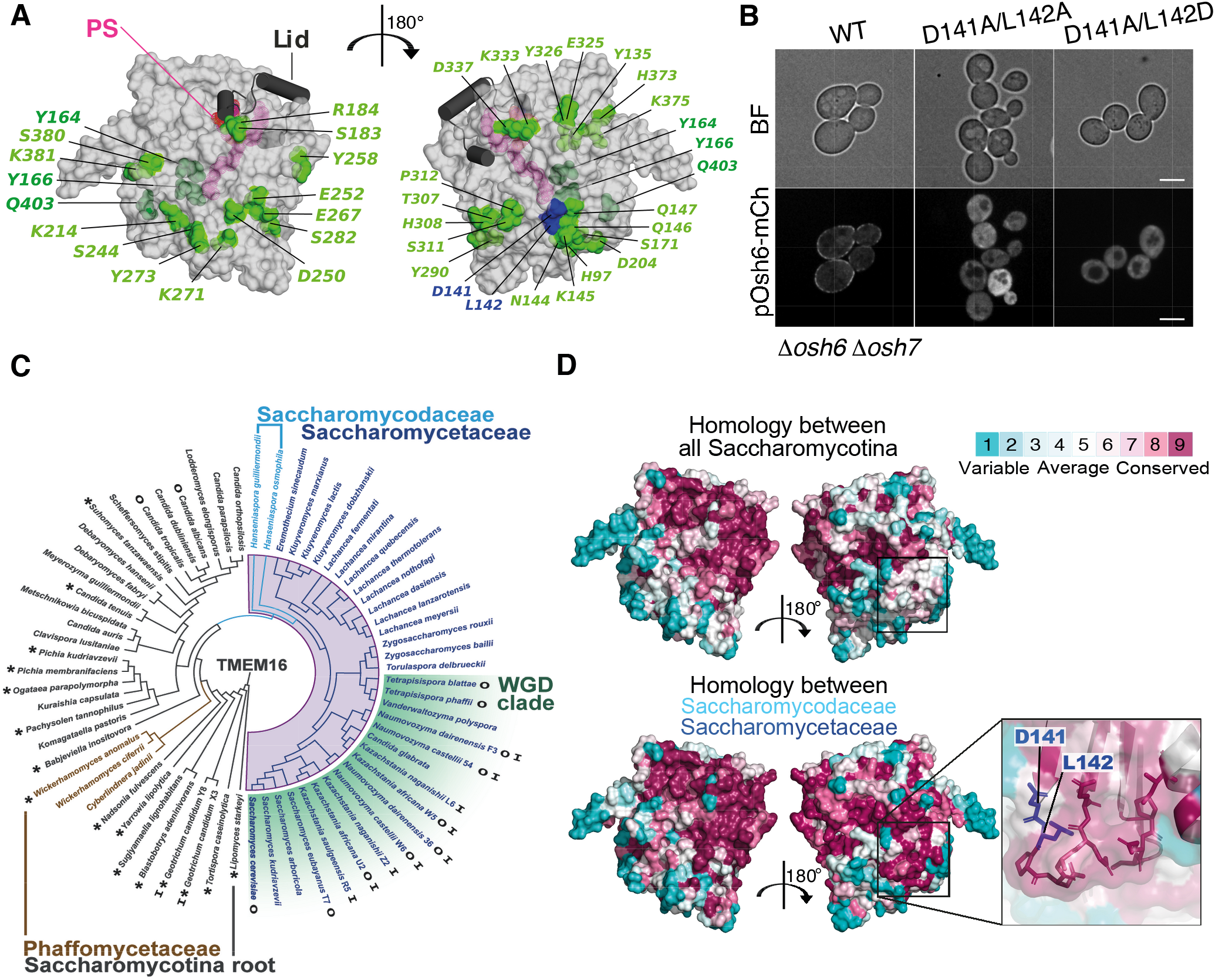
Identification of a conserved patch on Osh6 that interact with Ist2-tail. **(A)** Structure of Osh6 (PDB ID 4B2Z) showing the position of aa that were mutated (1 or 2 at a time) in the yeast two hybrid assay with Ist2 (aa 590-878) as bait. Substitution of aa in green did not affect the interaction, aa’s in blue affected the interaction. **(B)** Localization of Osh6-mCherry WT, Osh6^*L141A/D142A*^ and Osh6^*L141A/D142D*^ expressed from a plasmid in *osh6Δ osh7Δ* cells; representative images for several independent experiments are shown. **(C)** Evolutionary tree of TMEM16 homologs in Saccharomycotina. Species in blue (Saccharomycodaceae and Saccharomycetaceae monophyletic clades) contain orthologs with the conserved Osh6-binding region. The motif first appears in the Phaffomycetaceae paraphyletic group (brown color). Species in dark gray lack this motif. Asterisks indicate species with a short cytosolic tail (< 80 aa). WGD denotes whole genome duplication (green); ‘O’: two *OSH6* genes, ‘I’: two *IST2* genes. See also Fig. S7. **(D)** Osh6 surface representation showing the degree of aa conservation considering all sequences shown in (C) or only sequences from the Saccharomycetaceae and Saccharomycodaceae clades.

Osh6(D141A/L142A), fused to mCherry, was expressed at a similar level as WT Osh6-mCherry and was not thermally unstable (Fig. S6A). Full-length GFP-Ist2 extracted from yeast membranes co-immunoprecipitated WT Osh6-mCherry but not Osh6(D141A/L142A)-mCherry (Fig. 8A). Co-immunoprecipitation of Osh6-mCherry with Ist2 was specific because we could not detect any mCherry signal when Ist2 mutant with randomized linker (GFP-Ist2-RL) was used in the pull-down experiment under the same conditions. These results suggest that D141 and L142 residues of Osh6 comprise at least a part of the Ist2-binding site. However, analysis of protein localization suggested that the D141A/L142A substitution did not completely block the interaction between Osh6 and Ist2, as we could still detect some cortical Osh6(D141A/L142A)-mCherry in cells with a higher level of GFP-Ist2 when both proteins were expressed from a plasmid (Fig. S6B). In agreement, PS transport was blocked in yeast cells expressing Osh6(D141A/L142A) or Osh6(D141A/L142D) from a low-strength *CYC1* promoter, but PS was still transported to the PM when this mutant was expressed from the intermediate-strength *ADH1* promoter (*i.e.*, the promoter used in previous PS transport experiments) (Fig. 8B,C and Fig. S6C).

**Fig. 8.**
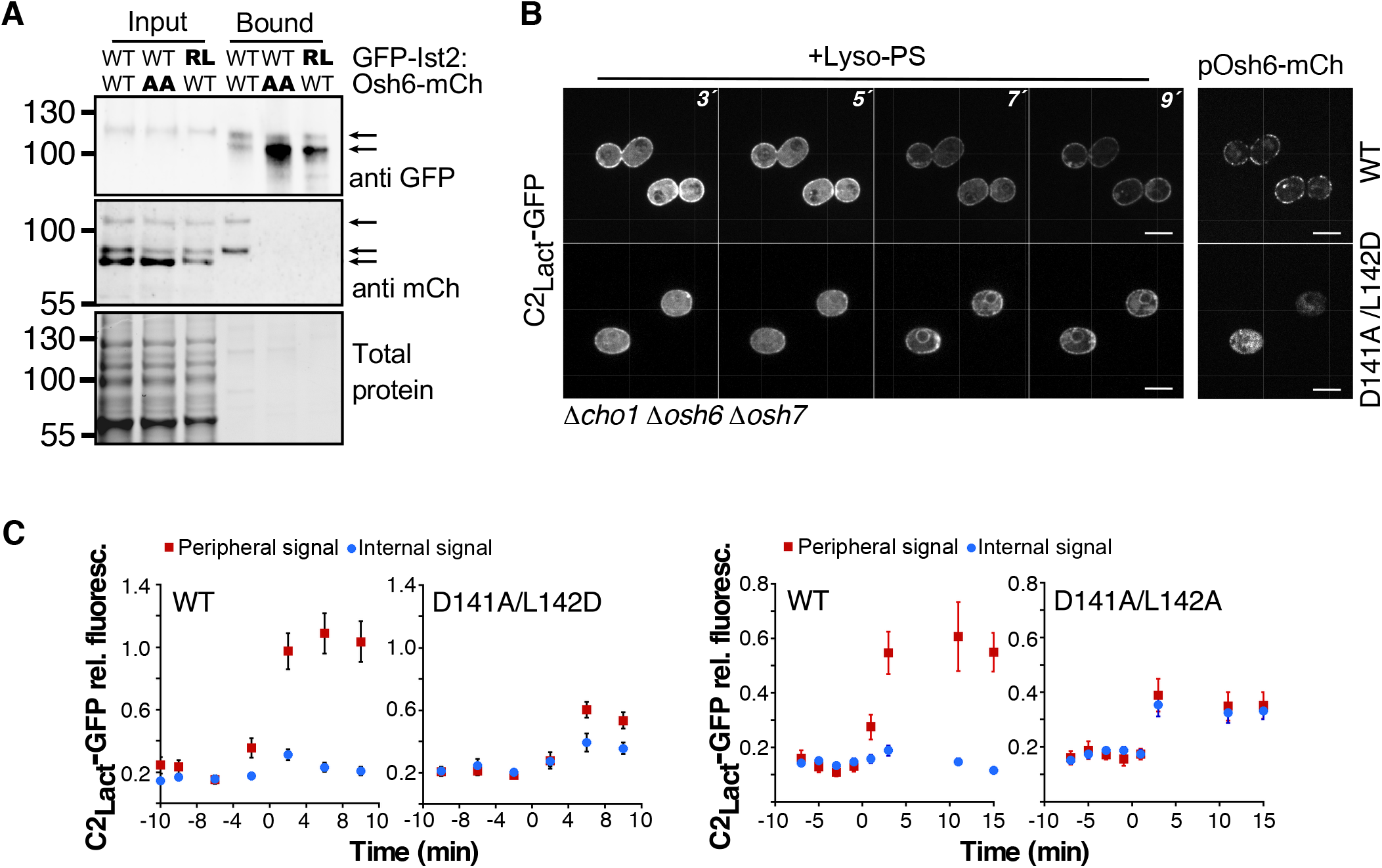
Osh6 D141/L142 mutants decrease interaction with Ist2 and PS transport. **(A)** Coimmunoprecipitation of GFP-Ist2 and Osh6-mCherry variants from *osh6Δ osh7Δ ist2Δ* cells. Lysates of cells expressing GFP-Ist2 (WT) or GFP-Ist2-RL (randomized linker) and Osh6-mCherry WT or D141A/L142A mutant (AA) were incubated with anti-GFP-magnetic beads, and analyzed by Western blot. For input, 0.3 cell OD equivalents were loaded for each sample; bound fractions are ~35x more concentrated than Input for α-mCherry detection, ~5x more concentrated for α-GFP detection. GFP-Ist2 and Osh-mCherry-specific bands are indicated with arrows. Note that some degradation of GFP-Ist2 (predicted size 135 kDa) likely occurred during purification. The predicted size of Osh6-mCherry is ~80 kDa; the two faster-migrating bands may differ in post-translational modification. The experiment was repeated >3 times. **(B)** Imaging of C2_Lact_-GFP over time in *cho1Δ osh6Δ osh7Δ* cells (depleted for PS) expressing Osh6-mCherry WT or D141A/L142D from low-level CYC1-promotor after addition of lyso-PS. **(C)** Quantification of C2_Lact_-GFP fluorescence peaks (peripheral and internal) over time in experiments with cells expressing Osh6-WT or D141A/L142D, or Osh6-WT or D141A/L142A, as shown in (B). Data are mean±s.e.m. (n=15) from one of two independent experiments. Scale Bar = 5 μm.

## Discussion

High PS concentration is a hallmark of the cytosolic leaflet of the PM and is important for diverse cellular processes, from establishment of cell polarity, signaling and cytoskeleton organization to control of endocytosis and caveolae formation (Fairn et al., 2011; Hirama et al., 2017; Nishimura et al., 2019; Sun and Drubin, 2013; Takano et al., 2008). Identification of proteins that specifically transport PS to the PM in yeast and in mammalian cells represented an important step towards understanding how this PS enrichment is achieved (Chung et al., 2015; Maeda et al., 2013; Moser Von Filseck et al., 2015). We now show that in yeast, the TMEM16 homologue Ist2, which acts as an ER-PM tether, is an obligatory partner for efficient transport of PS to the PM by Osh6, and likely also for its close paralog Osh7. Our analysis of fungal sequences suggests a co-evolution of Ist2 and Osh6 binding sites. Importantly, we demonstrate that even point mutations in the tail of Ist2 that block the interaction with Osh6 lead to a strong decrease in cellular PS levels, similar to *osh6Δ osh7Δ* deletion or to deletion of all tethering proteins in yeast or of the PI4 kinase IIIα in mammalian cells (Chung et al., 2015; Quon et al., 2018). These results suggest that the transport of PS from the ER to the PM by Osh6/7 is also important for preventing a build-up of PS in the ER membrane and therefore likely represents a major route for PS out of the ER.

In humans, the majority of ORP proteins contain additional targeting domains besides the ORD, with the exception of two truncated isoforms of ORP1 and ORP4, ORP1S and ORP4S (Raychaudhuri and Prinz, 2010). Few studies have looked at the function of these soluble isoforms (Charman et al., 2014; Jansen et al., 2011; Wang et al., 2019). In yeast, members of the family containing only an ORD are more common; besides Osh6 and Osh7, the related sterol/PI4P transporter Osh4 and its close paralog Osh5 also consist only of an ORD. Crystal structures of Osh4 and Osh6 show that these ORDs are quite large and, in addition to the core ß-barrel that encapsulates a lipid molecule, contain segments that could perform other functions (de Saint-Jean et al., 2011; Im et al., 2005; Maeda et al., 2013; Moser Von Filseck et al., 2015). We have identified a patch on the surface of Osh6 that participates in interaction with Ist2; this is the first time that an interaction between an ORD and another protein is clearly demonstrated. We predict that many more such interactions should exist in the cell to regulate the localization or function of ORP/Osh proteins. For example, a recent study has identified a phosphorylated loop in the ORD of ORP4 that regulates the association of ORP4 with intermediate filaments (Pietrangelo and Ridgway, 2019).

Why is it important to localize Osh6 to contact sites? Endogenously-tagged Osh6 can be observed both in the cytosol as well as at the cortical ER; however, our results show that its cortical localization, albeit at a low level, is necessary for lipid transport between the ER and the PM. Interestingly, we observed an accumulation of the PS reporter C2_Lact_-GFP at the ER when Osh6 was cytosolic due to disruption of its binding site on Ist2, suggesting that PS was not being extracted from the ER. It has been proposed that the main advantage of localizing LTPs to membrane contact sites is to increase the fidelity of lipid targeting (Hanada, 2018). However, our results point to the importance of LTP localization for efficient lipid transfer.

Only a small portion of the Ist2 tail is involved in binding to Osh6; what then is the function of the rest of the tail? Whereas its sequence is not conserved between fungal species, its length is generally high, far surpassing the length of an unfolded sequence that would be needed to traverse the width of ER-PM contacts. Such long tail could provide flexibility for the different steps of lipid transfer. The Ist2-interaction region on the Osh6 surface lies on the opposite side of the molecule than the entrance to the lipid binding pocket, suggesting that even when bound to Ist2, Osh6 could transiently interact with the ER and the PM membrane to extract/deliver its lipid ligands. The disordered Ist2 sequence could be important for regulating the dynamics of Osh6/7 within the crowded space of a membrane contact site, as has been recently shown for disordered sequences upstream of the ORDs of some mammalian ORPs (Jamecna et al., 2019).

Finally, it is intriguing that the membrane-embedded portion of Ist2 bears structural homology to the mammalian TMEM16 family of proteins, many of which have been shown to function as Ca^2+^-activated lipid scramblases and/or ion channels in the PM or possibly in the ER (Bushell et al., 2019; Falzone et al., 2018; Jha et al., 2019). PS is synthesized exclusively on the cytosolic side of the ER (Chauhan et al., 2016), but was proposed to be enriched in the luminal leaflet at steady state, although this was recently put under question (Fairn et al., 2011; Tsuji et al., 2019). Lipid scrambling would be another way to regulate the size of the PS pool available for transport by a cytosolic LTP. When reconstituted into liposomes, Ist2 did not display any lipid scramblase activity (Malvezzi et al., 2013). Nevertheless, it is tempting to speculate that there might be functional coupling between the ER-embedded portion of Ist2 and its tail that anchors Osh6 and enables PS transport.

## Materials and Methods

### Yeast Strains and Manipulations

Yeast manipulations were performed using standard methods and growth conditions. Yeast strains are listed in Table S2. Gene deletion and tagging were performed by homologous recombination and confirmed by PCR on genomic DNA. Yeast strains MdPY04 (*ist2*^736-743Δ^), MdPY07 (*cho1Δ ist2*^*736-743Δ*^), and MdPY06 (*ist2^T736A/T743A^*), were generated by CRISPR/Cpf1-mediated genomic editing (Swiat et al., 2017). Briefly, BY4742 or *cho1Δ* cells were transformed with pUDC175 coding the bacterial DNA-endonuclease fnCpf1 that mediates RNA-guided DNA cleavage at targeted genomic. Then, fnCpf1-carrying cells were transformed with pJMD_26 encoding a specific crRNA targeting *IST2* gene (CTTTACCAGAAACAATTCCAACATC), and a 100 bp homologous DNA fragment surrounding the target sequence for deletion of codons 736 to 743 (MdpY04 ad MdPY07) or to introduce mutations T736A and T743A (MdPY06). Cells were plated in synthetic dropout (SD) selective media lacking uracil and colonies were checked by PCR and sequencing. Before experiments, cells were grown in YPD until both plasmids (pJMD_26 and pUDC175) were lost.

### Plasmid construction

All plasmids used in this study are listed in Tables S3 and S4. QuikChange Site-Directed Mutagenesis Kit (Agilent) was used for generating point mutations in plasmids. Briefly, PCR amplification of plasmids using overlapping oligonucleotides carrying the desired mutation was performed according to the manufacturer’s instructions. I*n-Fusion* HD Cloning Plus kit (Takara Bio) was used, according to the supplier instructions, to generate plasmids: BFP-tagged Ist2-expressing plasmids pJMD_07, pJMD_08, pJMD_21; pJMD_01 (GFP-Ist2RL^729-747wt^) by replacement of codons 729 to 747 in the randomized linker of pAK76 with the corresponding *IST2*-WT sequence; similarly, pJMD_12 (BFP-IstRL^718-751wt^) was generated by replacing codons 718 to 751 in the randomized linker of pJMD_21 with the corresponding *IST2*-WT sequence. To generate plasmids used in the yeast two-hybrid assay (Table S4), *OSH4, OSH6* and *OSH7* ORFs and parts of the *IST2*-coding sequence were cloned into the BamHI and XhoI restrictions sites of pGADT7 or pGBKT7 using *Rapid DNA Ligation Kit* (Thermo Scientific). All plasmids were checked by DNA sequencing.

### Yeast Two-Hybrid Assay

For testing protein-protein interactions Matchmaker™ GAL4 Two-Hybrid System 3 was used according to the manufacturer’s instructions (Clontech). Briefly, yeast strain AH109 (James et al., 1996) was transformed with plasmids pGBKT7(*TRP1*) and pGADT7(*LEU2*) encoding the two proteins of interest fused to GAL4 DNA-Binding domain (bait) or GAL4 Activation domain (prey) (Table S4). Interaction between bait and prey initiates transcription of *HIS3* and *ADE2* (controlled by *GAL1* and *GAL2* promotor, respectively), allowing growth in restrictive media. Yeast cells were grown for 14-18 hours at 30°C, and 10-fold serial dilutions were plated on SD agar plates lacking leucine and tryptophan (control), leucine, tryptophan and histidine (No His), and leucine, tryptophan, histidine and adenine (No His/Ade). Cells were incubated for 3-5 days at 30°C before growth evaluation.

### Bimolecular Fluorescence Complementation (BiFc)

Osh6 and Osh7 were tagged with VN at their chromosomal loci in the BY4741 background using pFa6-VN::His3, and Ist2 was tagged in BY4742 with VC using pFa6-VC::kanMX (Sung et al., 2008). After mating and selection of the diploids on plates -Met -Lys, cells were grown to mid-logarithmic phase in SD medium at 30°C. Cells were mounted in the appropriate culture medium and imaged at room temperature with a motorized BX-61 fluorescence microscope (Olympus) equipped with a PlanApo 100× oil-immersion objective (1.40 NA; Olympus), a QiClick cooled monochrome camera (QImaging), and the MetaVue acquisition software (Molecular Devices). BiFc signals were visualized using a GFP filter set (41020 from Chroma Technology Corp.; excitation HQ480/20×, dichroic Q505LP, emission HQ535/50m). Alternatively (images shown in Fig. 1), cells were imaged using a spinning-disk confocal system, as described below.

### Fluorescence Microscopy

Yeast cells were grown 14-18 hours at 30°C in appropriate SD medium to maintain plasmid selection. When *cho1Δ, cho1Δ osh6Δ osh7Δ, cho1Δ* ist2^*736-743Δ*^ strains were grown, SD medium was supplemented with 1 mM ethanolamine. Cells were harvested by centrifugation in mid-logarithmic phase (OD_600_= 0.5–0.8) and prepared for viewing on glass slides when assessing protein localization at steady state. In the case of time-course experiments, visualization of C2_Lact_-GFP was performed using a microfluidics chamber (see below). When indicated, vacuolar staining was obtained by incubating cells with 100 μM CMAC (Life Technologies) for 10 min, after which the cells were washed twice before observation. Imaging was performed at room temperature, all images in this study (except BiFC shown in Fig. S2C, see above) were taken with an Axio Observer.Z1 microscope (Zeiss), equipped with an oil immersion plan apochromat 100x objective NA 1.4, a a sCMOS PRIME 95 camera (Photometrics), and a spinning-disk confocal system CSU-X1 (Yokogawa). GFP-tagged proteins, mCherry-tagged proteins and CMAC/BFP-tagged proteins staining were visualized with a GFP Filter 535AF45, RFP Filter 590DF35, and DAPI Filter 450QM60, respectively. BiFC signals were also visualized with the GFP filter. Images were acquired with MetaMorph 7 software (Molecular Devices). Images were processed with ImageJ (NIH) and with Canvas Draw (canvas X) for levels.

### Cellular PS transport assay

The cellular distribution of PS in cells lacking *CHO1* after the addition of exogenous lyso-PS was performed as described in (D’Ambrosio et al., 2019; Moser Von Filseck et al., 2015). Briefly, 18:1 lyso-PS (1-oleoyl-2-hydroxy-*sn*-glycero-3-phospho-L-serine, Avanti Polar Lipids), was dried under argon and resuspended in SD medium to 54 μM lyso-PS by vortexing and heating to 37°C. The lyso-PS suspension was always prepared fresh, maintained at room temperature and used within 1-3 hours after preparation. PS transport assay was carried out using a Microfluidic Perfusion Platform (ONIX) driven by the interface software ONIX-FG-SW (Millipore). Strains *cho1Δ, cho1Δ osh6Δ osh7Δ* and *cho1Δ* ist2 ^*736-743Δ*^, transformed with pC2_Lact_-GFP and other plasmids, as indicated, were injected into a YO4C microfluidics chamber and maintained in a uniform focal plane. We always imaged two strains simultaneously to control for small differences between experiments due to variability in the stability of the lyso-PS suspension, which is difficult to control (D’Ambrosio et al., 2019). For this, we stained one of the two strains with CMAC as described above just before imaging, then mixed the strains at a ratio 1:1 before injecting them in the chamber. Normal growth conditions were maintained by flowing cells with SD medium or SD medium containing lyso-PS at 3 psi. Cells were imaged every 2 min over a total time of 30 to 40 min, starting when lyso-PS-containing medium was injected into the system. Cells were imaged in 5 z-sections separated by 0.7 μm, afterwards manually selecting for the best focal plane, in order to correct for any focal drift during the experiment. Usually we imaged cells in four different fields of 840 x 1140 pixels.

### Image Analysis

All image analysis was done using Image-J (NIH). Quantification of Osh6 distribution between cortical and cytosolic only was performed manually, by counting the cells in which some enrichment of cortical Osh6 fluorescence could be observed, versus cells where Osh6 signal was uniformly cytosolic. Quantification of Osh6-mCherry distribution as a function of GFP-Ist2 or BFP-Ist2 fluorescent signal was performed by profiling cell signal intensity across two transversals line that were manually placed on a single z-section of each cell in the Ist2-fluorescence channel. Ist2 fluorescence was calculate as the mean of four peripheral fluorescence peaks. The same lines were used to measure peripheral fluorescence of Osh6-mCherry in the red channel, divided by the average internal (cytosolic) fluorescence after subtraction of background fluorescence.

Steady-state distribution of C2_Lact_-GFP was analyzed on a single z-section of each cell. Using wand (tracing) tool, the external limit of the cell (perimeter) was selected and total cell fluorescence was measured (*Analyze/measure*). Subsequently, internal fluorescence was measure (*Analyze/measure*) after reducing the cell perimeter (*Edit/selection/enlarge, pixels=−7*). Peripheral fluorescence (difference between total and internal fluorescence) was normalized to total fluorescence and plotted like “peripheral signal”.

Quantification of peripheral peaks (mostly PM and cER signal) and internal (mostly perinuclear ER) was performed by profiling cell signal intensity across a transversal line drawn in the cell. Intensities of the peaks are quantified and normalized relative to the total signal. The cell profile (peripheral and internal peaks) is followed and quantified over time of the experiment. Time = 0 is set as the point in which C2_Lact_-GFP distribution in WT cells starts changing (moving from cytosolic to internal and peripheral localization).

Data were processed in Excel and plotted using Kaleidagraph 4.5 (Synergy Software). We carried out the statistical analysis using KaleidaGraph. To compare the means of multiple groups, we used one-way ANOVA followed by Tukey’s multiple comparisons.

### Preparation of yeast protein extracts

Yeast cultures used were grown over-night in appropriate SD medium to mid-logarithmic phase. 1 OD_600_ of yeast cells was with TCA (10% final concentration) for 10 min on ice to precipitate proteins, centrifuged at 16,000g for 10 min at 4°C, and mechanically disrupted with glass beads using a vortex for 10 min at 4°C. Lysates were transferred to a fresh tube and centrifuged again. The protein pellets were washed twice with ice-cold acetone, dried and resuspended in 50 μl of sample buffer (50 mM Tris-HCl, pH 6.8, 100 mM DTT, 2% SDS, 0.1% bromophenol blue, 10% glycerol), complemented with 50 mM Tris-base. Protein samples were heated at 55°C for 10 min and 10 μl was loaded on SDS–PAGE (4-20% Mini-PROTEAN TGX Stain-Free, Bio-Rad) and analyzed by Western blot.

### Immunoprecipitation of Ist2-GFP

To check interaction between Osh6-mCherry and GFP-Ist2 in cells, we immunoprecipitated GFP-Ist2 from strain ACY406 (*ist2Δ osh6Δ osh7Δ*) transformed with pAK75 (GFP-Ist2) and pOsh6-mCherry, or plasmids expressing GFP-Ist2 and Osh6-mCherry mutants. 60 OD_600_ equivalents of yeast cells were grown to mid-logarithmic phase in minimum medium at 30°C, washed with distilled water and snap-frozen in liquid nitrogen. Yeast pellets were resuspended in 2 x 500 μl of ice-cold lysis buffer (50mM Hepes, 150mM NaCl, 1% NP-40, 1mM EDTA) containing protease inhibitors (complete Protease Inhibitor Cocktail, Roche Life Sciences), 2 mM phenylmethylsulfonyl fluoride (Sigma-Aldrich) and phosphatase inhibitors (PhosSTOP, Roche). Cells were mechanically disrupted with glass beads using a vortex for 8 min at 4°C. The lysates were centrifuged at 3000g for 5 min at 4°C and the supernatants were collected. 50 μl of each supernatant was diluted to 1 ml and precipitated with 10% TCA (for Input); the rest was incubated with 15 μl of GFP-Trap_MA magnetic agarose beads (ChromoTek GmbH, Planegg-Martinsried, Germany), prewashed with lysis buffer, for 2.5 hours at 4°C. The beads were collected using a magnetic rack and washed three times in 1 mL of lysis buffer; finally, they were resuspended in 50 μl of sample buffer (50 mM Tris-HCl, pH 6.8, 6% SDS, 0.01% bromophenol blue) and then incubated for 10 min at 37°C. TCA precipitated samples were incubated on ice for >10 min and centrifuged at 16,000g for 10 min at 4°C. The pellets were washed 2 x with 500 μl of ice-cold acetone, resuspended in 50 μl of sample buffer with urea (50 mM Tris-HCl, pH 6.8, 100 mM DTT, 3% SDS, 3M urea, 0.1% bromophenol blue, 10% glycerol) complemented with 50 mM Tris-base, vortexed and incubated for 10 min at 37°C. Samples (10 μl for input and bound fractions for mCherry detection, 1.5 μl for bound franction for GFP detection) were loaded on SDS–PAGE (4-20% Mini-PROTEAN TGX Stain-Free, Bio-Rad) and analyzed by Western blot.

### Western blot analysis

After electrophoresis, total proteins were visualized in the TGX Stain-Free gels (Bio-Rad) after 1 min UV-induced photoactivation with Gel Doc EZ Imager (Bio-Rad). Proteins were then transferred onto a nitrocellulose membrane. GFP-Ist2 was detected with a rabbit polyclonal anti-GFP antibody (ThermoFisher ref. A11122, 1:5000 dilution), and Osh6-mCherry was detected with a rabbit polyclonal anti-RFP antibody (ThermoFisher ref. 10041338, 1:1500 dilution). Horseradish peroxidase–coupled secondary antibodies were from Sigma (anti-mouse, ref A5278; anti-rabbit, ref. A6154; both used at 1:5000 dilution). Chemiluminescence signals were acquired with Gel Doc EZ Imager.

### Osh6-TAP purification

Osh6 was tagged with a TAP epitope at its chromosomal locus in the BY4741 background using the plasmid pYM13 (Janke et al., 2004). Two liters of BY4741 and Osh6-TAP cells were grown overnight at 30°C to an OD_600_ of 0.7. The cultures were centrifuged at 4000 rpm for 5 minutes.

Cell pellets were washed once with cold sterile water and resuspended in 4.5 ml of lysis buffer (50 mM Tris-HCl, pH 8.0, 100 mM NaCl, 5mM MgCl2, 1% Triton X-100, 1mM PMSF, and complete protease inhibitor cocktail; Roche) and lysed twice with high pressure (1200 psi) at −80 °C using a cell breaker (Carver, Inc.). Lysates were cleared by centrifugation and the cleared lysates were incubated with 30 μl of IgG-coupled magnetic beads (Dynabeads; Thermo Fisher Scientific) for 1h at 4°C. Beads were collected with a magnetic rack and washed three times in 1 ml lysis buffer containing 0.5% Triton X-100 and then 2 times in the same buffer without detergent. Beads were then resuspended in 400 μl of lysis buffer and 10 μl of AcTEV (Invitrogen) was added and incubated at 18°C overnight. Beads were discarded with a magnet, eluted proteins were divided in two tubes and precipitated using chloroform-methanol. One pellet was resuspended in 15 μl sample buffer, heated at 95°C for 5 min and loaded on a NuPAGE 4–12% gradient polyacrylamide gel. The other pellet was used for the mass spectrometry.

### Proteomic Analysis by Mass Spectrometry

Proteins were digested overnight at 37°C in 20 ml of 25 mM NH_4_HCO_3_ containing sequencing-grade trypsin (12.5 mg/ml; Promega). The resulting peptides were sequentially extracted with 70% acetonitrile and 0.1% formic acid. Digested samples were acidified with 0.1% formic acid. All digests were analyzed by an Orbitrap Fusion equipped with an EASY-Spray nanoelectrospray ion source and coupled to an Easy nano-LC Proxeon 1000 system (all from ThermoFisher). Chromatographic separation of peptides was performed with the following parameters: Acclaim PepMap100 C18 precolumn [2 cm, 75 mm inner diameter (i.d.), 3 mm, 100 Å], Pepmap-RSLC Proxeon C18 column [50 cm, 75 mm i.d., 2 mm, 100 Å], 300 nl/min flow, using a gradient rising from 95% solvent A (water, 0.1% formic acid) to 40% B (80% acetonitrile, 0.1% formic acid) in 120 min, followed by a column regeneration of 20 min, for a total run of 140 min. Peptides were analyzed in the Orbitrap in full-ion scan mode at a resolution of 120,000 [at m/z (mass/charge ratio) 200] and with a mass range of m/z 350 to 1550 and an AGC target of 2 × 105. Fragments were obtained by higher-energy C-trap dissociation activation with a collisional energy of 30% and a quadrupole isolation window of 1.6 Da. MS/MS data were acquired in the linear ion trap in a data-dependent mode, in top-speed mode with a total cycle of 3 s, with a dynamic exclusion of 50 s and an exclusion duration of 60 s. The maximum ion accumulation times were set to 250 ms for MS acquisition and 30 ms for MS/MS acquisition in parallelization mode. Data were processed with Proteome Discoverer 1.4 software (ThermoFisher) coupled to an in-house Mascot search server (Matrix Science; version 2.4). The mass tolerance of fragment ions was set to 7 ppm for precursor ions and 0.5 Da for fragments. Identification of tryptic peptides related to proteins were performed on *Saccharomyces cerevisiae* taxonomy from Swissprot database. Q-values of peptides were calculated using the percolator algorithm, and a 1% filter was applied as a false-discovery rate threshold.

### Lipidomic Analysis by Mass Spectrometry

Sample preparation was carried out according to (Klose et al., 2012). In short, BY4247, *osh6Δosh7Δ*, *ist2Δ*, *ist2*^736-743Δ^, *ist2*^T736A/T743A^ cells were grown in triplicates in Erlenmeyer flasks (180 rpm, 30°C) until OD_600_ ~0.5 was reached. 10 ODs of cell were centrifugated (3min, 5000 g) and washed twice with 155 mM ammonium bicarbonate. Supernatant was discarded and pellets were snap-frozen in liquid nitrogen and kept in −80°C until lipid extraction.

Lipids were extracted according to a modified Bligh and Dyer protocol. The yeast pellet was collected in a 1.5 mL Eppendorf tube and 200 μL of water was added. After vortexing (30s), the sample was transferred to a glass tube containing 500 μL of methanol and 250 μL of chloroform. The mixture was vortexed for 30s and centrifuged (2500 rpm, 4°C, 10 min). Then 300 μL of the organic phase was collected in a new glass tube and dried under a stream of nitrogen. The dried extract was resuspended in 60 μL of methanol/chloroform 1:1 (v/v) and transferred in an injection vial.

Reverse phase liquid chromatography was selected for separation with an UPLC system (Ultimate 3000, ThermoFisher). Lipid extracts were separated on an Accucore C18 150×2.1, 2.5 μm column (ThermoFisher) operated at 400 μl/min flow rate. The injection volume was 3 μl. Eluent solutions were ACN/H_2_O 50/50 (V/V) containing 10mM ammonium formate and 0.1% formic acid (solvent A) and IPA/ACN/H_2_O 88/10/2 (V/V) containing 2mM ammonium formate and 0.02% formic acid (solvent B). The step gradient of elution was in %B : 0.0 min, 35%; 4.0 min, 60 %; 8.0 min, 70%; 16.0 min, 85%; 25.0 min, 97%. The UPLC system was coupled with a Q-exactive Mass Spectrometer (ThermoFisher, CA); equipped with a heated electrospray ionization (HESI) probe. This spectrometer was controlled by Xcalibur software and operated in electrospray positive mode. Data were acquired with dd-MS2 mode at a resolution of 70 000 for MS and 35 000 for MS2 (200 m/z) and a normalized collision energy (NCE) of 25 and 30 eV. Data were reprocessed using Lipid Search 4.1.16 (ThermoFisher). The product search mode was used and the identification was based on the accurate mass of precursor ions and MS2 spectral pattern.

### Sequence alignment and phylogeny of Ist2

A first set of sequences from the Saccharomycotina subphylum (Tax ID: 147537) and corresponding to TMEM16 homologues (IPR007632) was retrieved from Interpro database (Finn et al., 2017). We obtained 68 sequences belonging to 62 species. Sequences were aligned using MAFFT (E-INS-I) algorithm (Katoh et al., 2019) with the default parameters of Jalview program (v2.11.0) (Waterhouse et al., 2009). One duplicated homologue from *Kazachstania saulgeensis* was deleted (see Fig. S7). A phylogenetic inference was done using the maximum likelihood method PhyML (-LG matrix, aLRT (SH-like) branch support, BIONJ and NNI for tree searching) (Guindon et al., 2010). The tree was displayed and formatted on FigTree v1.4.4 and *Lipomyces starkeyi* was selected as the rooting taxa. In order to display the highly conserved region that interacts with Osh6/Osh7, sequences were ordered on Jalview according to the phylogeny of Saccharomycotina (Shen et al., 2018), and we deleted sequences that do not belong to the monophyletic group including Saccharomycetaceae and Saccharomycodaceae, as indicated by our phylogenetic tree. The alignment conservation scores for each residue position of Ist2 were extracted from this subset and were displayed as a heatmap using Prism 8.3 (GraphPad Software). See Fig. S7 for the full alignment.

### Sequence alignment and structural analysis of Osh6

A second set of Osh6/Osh7 homolog sequences (i.e., containing the LPTFILE motif, as previously described; (Maeda et al., 2013) were retrieved from the same 58 Saccharomycotina species. The resulting 70 sequences of Osh6/Osh7 homologs were aligned using MAFFT-G-INS-I with BLOSUM 80 matrix. Alignments were subsequently analyzed with Jalview. Amino-acid evolution rate was calculated using Consurf upon 4PH7_D PDB structure and using maximum likelihood method with -LG Matrix (Ashkenazy et al., 2016). Consurf results were displayed with Pymol software (Version 2.0 Schrödinger, LLC).

## Acknowledgments

We thank E Sajovic, A Dubois and G Žun for help with experiments, and A-C Gavin, CL Jackson, S Leon, U Petrovič and L Veenhoff for strains and plasmids. We acknowledge the IJM ImagoSeine facility, member of IBiSA and the France-BioImaging infrastructure (ANR-10-INBS-04), the IJM Proteomics facility [supported by the Region Ile-de-France (SESAME), the Paris-Diderot University (ARS), and CNRS], and B Antonny, and S Leon and members of Jackson-Verbavatz team for helpful discussions.

## Author Contributions

JM D’Ambrosio, V Albanèse and A Čopič performed experiments. NF Lipp performed in silico analyses with G Drin. L Fleuriot and D Debayle performed lipidomic analysis. JM D’Ambrosio, V Albanèse, NF Lipp, G Drin and A Čopič designed research and analyzed data. A Copic conceived the study. A Copic and JM D’Ambrosio wrote the manuscript, with input from all other authors.

