## Supplemental Figures and Tables for "Osh6 requires Ist2 for localization to the ER-PM contacts and efficient phosphatidylserine transport"

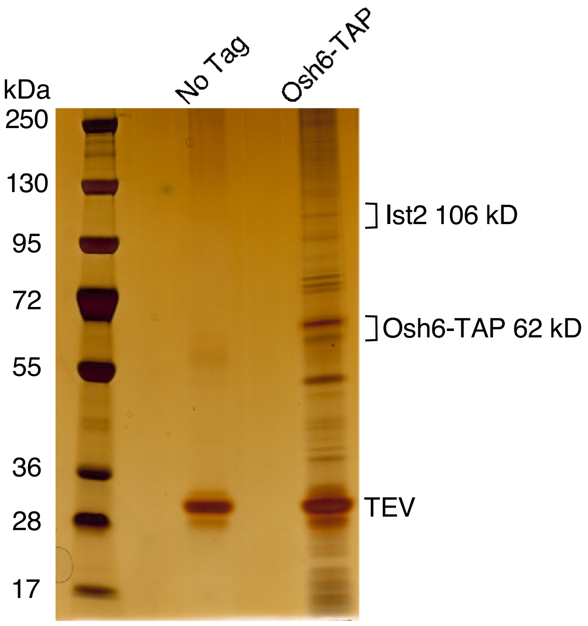


***Fig. S1. Purification of Osh6-TAP from WT cells.*** *After elution with TEV protease, eluates were loaded on SDS-PAGE and the gel was silver stained. Approximate positions of Ist2, Osh6-TAP and TEV are indicated. Experiment was performed once.*

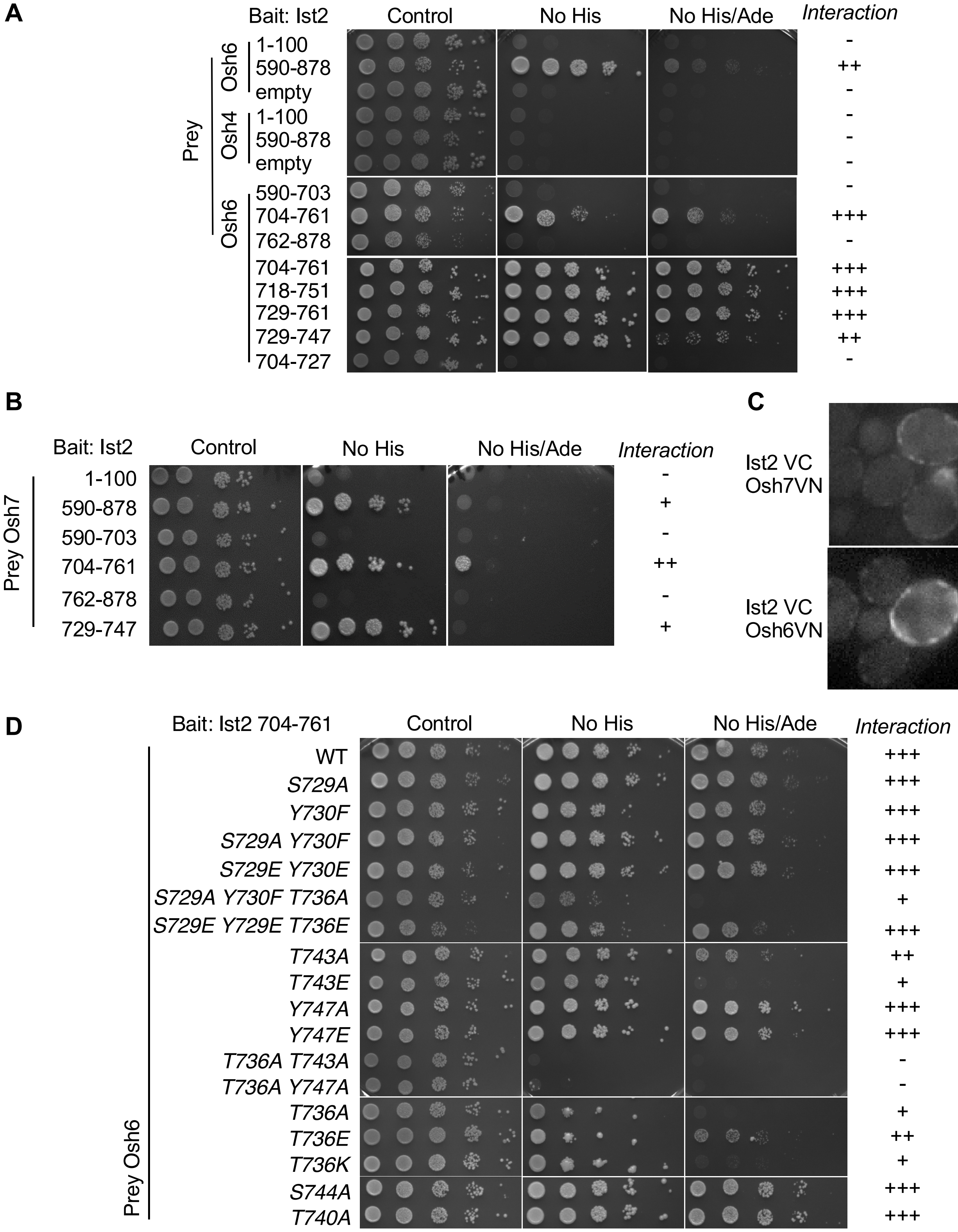


***Fig. S2. Yeast two-hybrid analysis of Ist2 interaction with Osh6 and Osh7. (A)*** *10-fold serial dilutions of yeast cells expressing Ist2 fragments (first and last aa are indicated) fused to GAL4 DNA-binding domain (bait) and full-length Osh6 or Osh4 fused to GAL4 Activation domain (prey) on control, SD-His and SD-His-Ade reporter plates. Plates were incubated at 30°C for 3 days.* ***(B)*** *10-fold serial dilutions of yeast cells expressing Ist2 fragments fused to GAL4 DNA-Binding domain (bait) and full-length Osh7 fused to GAL4 Activation domain (prey) on control, and SD-His reporter plate. Plates were incubated at 30°C for 3 days.* ***(C)*** *BiFC in diploid cells expressing endogenously tagged Osh7-VN and Ist2-VC or Osh6-VN and Ist2-VC, as indicated. Scale Bar = 5 µm.* ***(D)*** *10-fold serial dilutions of yeast cells expressing Ist2 fragment aa704-761 fused to GAL4 DNA-Binding domain (bait) and different Osh6 mutants fused to GAL4 Activation domain (prey) on control and SD-His and SD-His-Ade reporter plates. Plates were incubated at 30°C for 3 days. Interaction score: (-) No growth detected in -His, (+) growth detected in -His, (++) weak growth detected in -His/-Ade, (+++) Strong growth detected in -His/-Ade. All plates are representative of 3 independents experiments.*

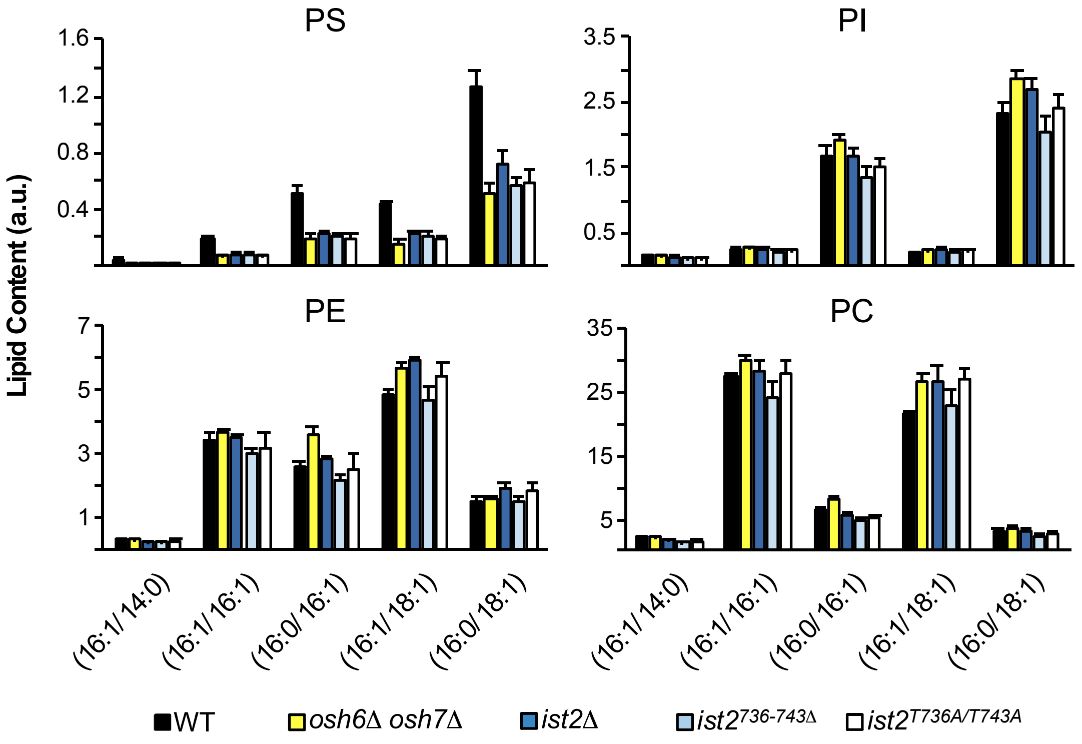


**Fig. S3. Lipidomic analysis of total cellular content of PC, PE, PI and PS species in WT, *osh6*Δ *osh7*Δ and *ist2* mutant cells.** Lipids were extracted from WT, *osh6*Δ *osh7*Δ, *ist2*Δ, *ist2*$^{736\text{-}743\Delta}$ and *ist2*$^{T736A/T743A}$ (chromosomal deletion of 8 codons) and *ist2*$^{T736A/T743A}$ (chromosomal substitution in 2 codons) cells, and analyzed by mass spectrometry. Data are mean±s.d. from 3 independent samples. Parentheses denote acyl chain length and saturation. The experiment was performed 3 times, yielding similar results.

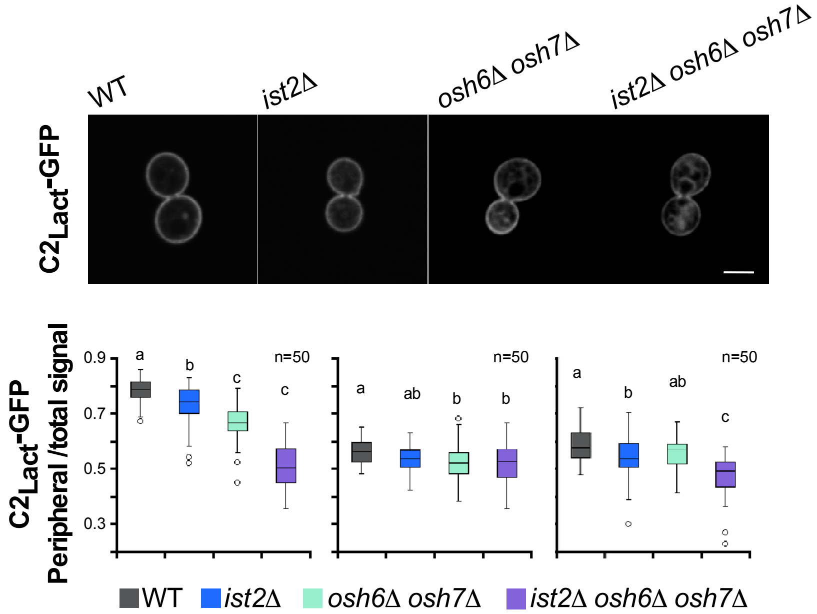


**Fig. S4. Steady-state distribution of PS in WT, *ist2*Δ and *osh6*Δ *osh7*Δ cells.** Subcellular localization of $C2_{lact}$-GFP in WT, *ist2*Δ, *osh6*Δ *osh7*Δ and *osh6*Δ *osh7*Δ *ist2*Δ cells was assessed by fluorescence microscopy (top panels). Quantification of relative $C2_{Lact}$-GFP peripheral signal, normalized to total cellular fluorescence, is shown in box plots for 50 individual cells (n=50) in 3 independents experiments. Each box encloses 50% of the data, median value is displayed as a line, top and bottom of the box mark the limits of ± 25%. The lines extending from the top and bottom of each box mark the minimum and maximum values. Outlier are displayed as an individual point. Different letters (“a”, “b” and “c”) indicate significant differences between the means, “ab” denotes non-significant difference from either mean “a” or mean “b” (multiple comparison using one-way ANOVA, posthoc Tukey’s test at $p < 0.05$).

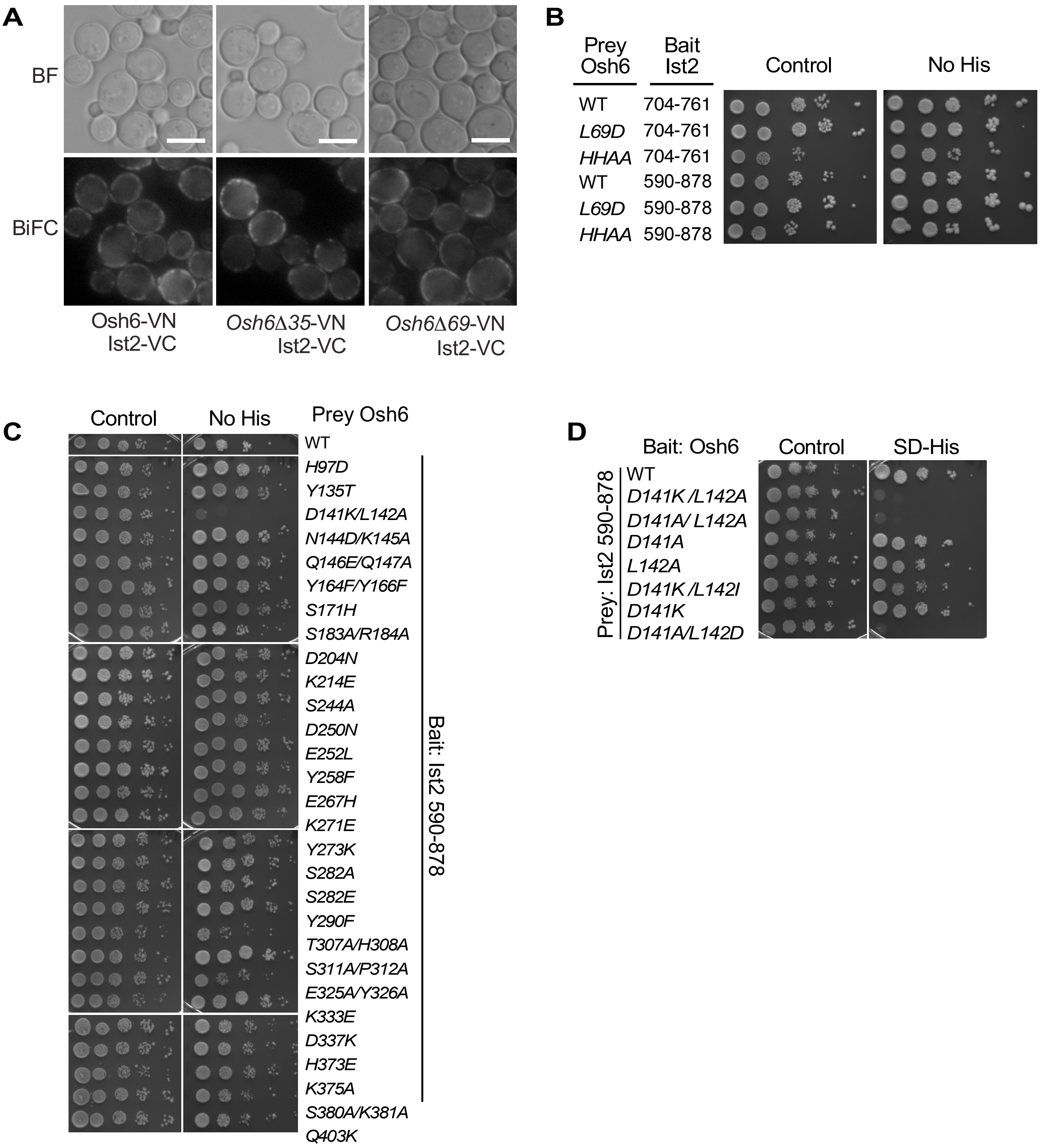


**Fig. S5. Mapping Ist2 interaction site on Osh6. (A)** Bimolecular fluorescence complementation (BiFC) between Osh6 WT, or two N-term deletion mutants Osh6Δ35 (lacking 35 N-term aa, which are disordered) or Osh6Δ69 (lacking the whole N-term lid that covers the lipid-binding pocket) fused to Venus N-term, and Ist2 fused to Venus C-term. Scale Bar= 5 µm. **(B-D)** 10-fold serial dilutions of yeast cells (AH109) expressing Ist2 full-length cytosolic tail (aa 590-878) and Osh6 WT or selected mutants with substitutions in indicated aa, fused to GAL4 Activation domain (prey) or GAL4 DNA-Binding domain (bait), respectively. Osh6 mutants were selected based on structural and conservation analysis. Interaction of bait and prey proteins activates transcription of *HIS3* (controlled by GAL promoter) and allows growth in SD-His selective media. Results are representative of 3 independents experiments.

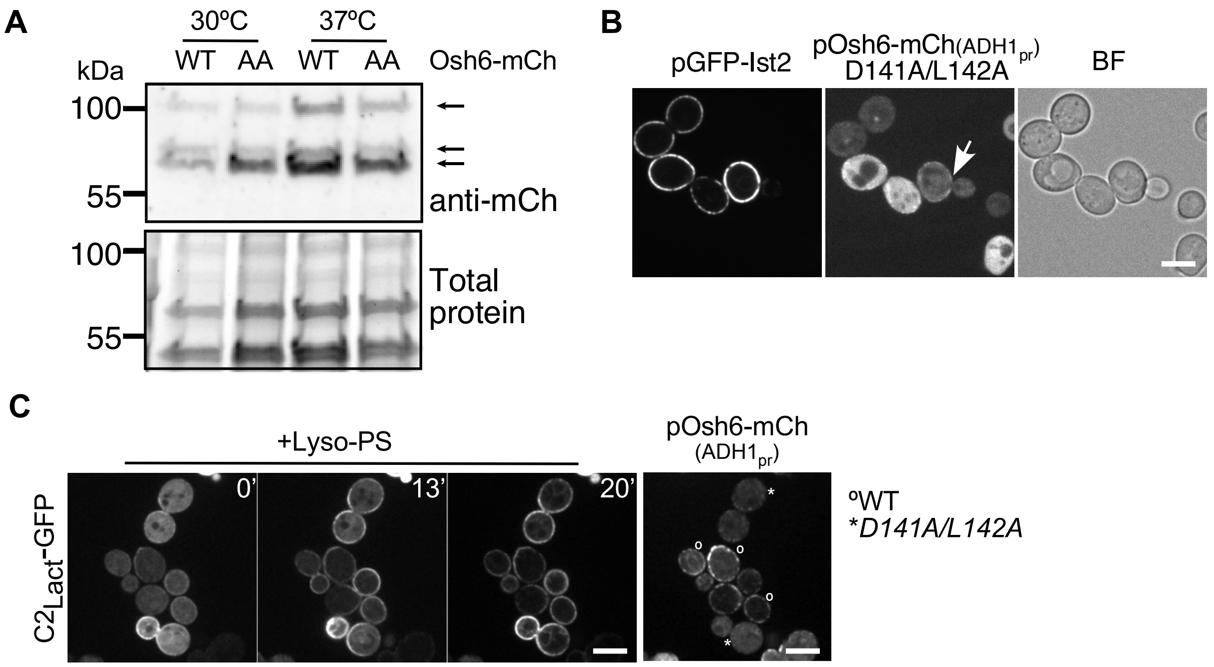


**Fig. S6. Mutations of D141 and L142 do not affect the stability of Osh6 and still permit some interaction with Ist2. (A)** Western blot analysis of whole cell TCA precipitates showing mCherry-tagged Osh6 and Osh6$^{D141A/L142A}$ protein levels at 30$^{O}$C and 37$^{O}$C. Experiment was repeated 2 times. **(B)** GFP-Ist2 and Osh6$^{L141A/D142A}$ -mCherry, expressed from low-copy plasmids in *ist2*Δ *osh6*Δ *osh7*Δ cells were visualized using fluorescence microscopy. **(C)** Imaging of C2$_{Lact}$-GFP over time in a mix of two *cho1*Δ *osh6*Δ *osh7*Δ strains (lacking endogenous PS), expressing Osh6-mCherry WT ($^{O}$) or D141A/L142D mutant (*) from the medium-level ADH1-promotor. For identification, cells expressing WT Osh6 were labelled with CMAC. Time (in min) after addition of lyso-PS is indicated. Osh6-mCherry signal is shown in the right panel at t=0 min. Representative images from two independent experiments are shown. Scale Bar = 5 µm.

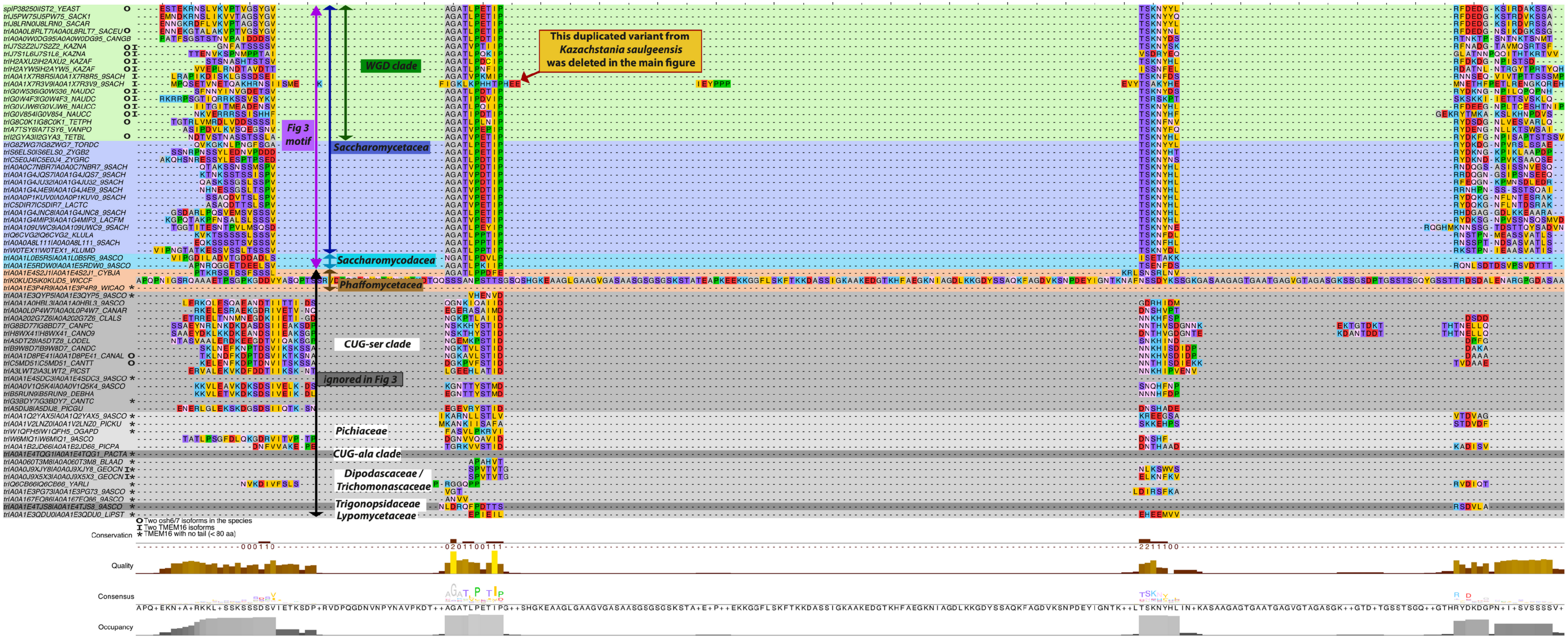

This duplicated variant from Kazachstania saulgeensis was deleted in the main figure
WGD clade
Fig 3 motif
Saccharomycetacea
Saccharomycodacea
Phaffomycetacea
CUG-ser clade
ignored in Fig 3
Pichiaceae
CUG-ala clade
Dipodascaceae / Trichomonascaceae
Trigonopsidaceae
Lypomycetaceae
Two osh6/7 isoforms in the species
Two TMEM16 isoforms
TMEM16 with no tail (< 80 aa)
Conservation
Quality
Consensus
Occupancy


**Figure S7**

***Fig. S7. Related to Fig. 3 and Fig. 7C: Alignment of the tail region of 68 TMEM16 homologous sequences from 62 Saccharomycotina species.*** *Color coding is explained in the figure and is the same as in Fig. 7C. Conservation score and heatmap on a brown to yellow scale are shown at the bottom. Asterisks indicate homologues with a short cytosolic tail (< 80 aa). WGD denotes whole genome duplication (green); 'O': two OSH6 genes, 'I': two IST2 genes.*

**Table S2.** Yeast strains used in this study.

| Name | Alias | Background | *Genotype* | Origin/ Reference |
|---|---|---|---|---|
| BY4742 | WT | BY4742 | *MATα his3Δ1 leu2Δ0 lys2Δ0 ura3Δ0* | *Euroscarf* |
| BY4741 | *WT* | | *Mata his3Δ1 leu2Δ0 met15Δ0 ura3Δ0* | *Euroscarf* |
| VAY2817 | *OSH6*-TAP | BY4742 | *MATα his3Δ1 leu2Δ0 lys2Δ0 ura3Δ0 OSH6-TAP::KanMX* | *This study* |
| Ist2-GFP | *IST2-GFP* | BY4741 | *MATa his3Δ1 leu2Δ0 met15Δ0 ura3Δ0 IST2-GFP::HIS3* | *Huh et al., 2003* |
| Osh6-GFP | *OSH6-GFP* | BY4741 | *MATa his3Δ1 leu2Δ0 met15Δ0 ura3Δ0 OSH6-GFP::HIS3* | *Huh et al., 2003* |
| VAY2802 | *OSH6-GFP ist2Δ* | BY4741 | *MATα his3Δ1 leu2Δ0 lys2Δ0 ura3Δ0 ist2Δ::hphNT1 OSH6-GFP::KanMX* | *This study* |
| VAY3652 | *OSH6-GFP scs2Δ scs22Δ* | BY | *MAT ? his3Δ1 leu2Δ0 scs2Δ::KanMX Δscs22::KanMX Osch6-GFP::HIS3* | *This study* |
| VAY2811 | *IST2-VC* | BY4742 | *MATα his3Δ1 leu2Δ0 lys2Δ0 ura3Δ0 IST2-VC::KanMX* | *This study* |
| VAY2695 | $ist2^{\Delta590}$-VC | BY4742 | *MATα his3Δ1 leu2Δ0 lys2Δ0 ura3Δ0* $IST2^{\Delta590}$-*VC::KanMX* | *This study* |
| VAY2829 | *OSH6-VN* | BY4741 | *MATa his3Δ1 leu2Δ0 met15Δ0 ura3Δ0 OSH6-VN::HIS3* | *This study* |
| VAY2969 | *OSH7-VN* | BY4741 | *MATa his3Δ1 leu2Δ0 met15Δ0 ura3Δ0 OSH7-VN::HIS3* | *This study* |
| AH109 | AH109 | AH109 | *MATa trp1-901 leu2-3,112 ura3-52 his3-200 gal4Δ gal80Δ LYS2::*$GAL1_{UAS}$-$GAL1_{TATA}$-*HIS3, *$GAL2_{UAS}$-$GAL2_{TATA}$-*ADE2 URA3::*$MEL1_{UAS}$-$MEL1_{TATA}$-*lacZ* | *James et al.,1996* |
| VAY2784 | *ist2Δ* | BY4742 | *MATα his3Δ1 leu2Δ0 lys2Δ0 ura3Δ0 ist2Δ::hphNT1* | *This study* |
| MdPY08 | *cho1Δ* | BY4742 | *MATα his3Δ1 leu2Δ0 lys2Δ0 ura3Δ0 cho1Δ::kanMX* | *This study* |
| SGAY1635 | *osh6Δ osh7Δ* | BY4742 | *MATα can1Δ::STE2pr-LEU2 lyp1Δ ura3Δ0 leu2Δ0 his3Δ1 met15Δ0 osh6Δ::HYGMX osh7Δ::natNT2* | *Maeda et al., 2013* |
| SGAY7039 | *cho1Δ osh6Δ osh7Δ* | BY4742 | *MATα can1Δ::STE2pr-LEU2 lyp1Δ ura3Δ0 leu2Δ0 his3Δ1 met15Δ0 cho1Δ::KanMX osh6Δ::hphNT1 osh7Δ::natNT2* | *Maeda et al., 2013* |
| ACY406 | *ist2Δ osh6Δ osh7Δ* | SGAY1635 | *MATa can1Δ::STE2pr-LEU2 lyp1Δ ura3Δ0 leu2Δ0 his3Δ1 met15Δ0 ist2Δ::HISMX6 osh6Δ::hphNT1 osh7Δ::natNT2* | *This study* |
| MdPY01 | *ist2Δ osh6Δ osh7Δ* | ACY406 | *MATa can1Δ::STE2pr-LEU2 lyp1Δ ura3Δ0 leu2Δ0 his3Δ1 met15Δ0 ist2Δ::kanMX osh6Δ::hphNT1 osh7Δ::natNT2* | *This study* |
| MdPY04 | $ist2^{736\text{-}743\Delta}$ | BY4742 | *MATα his3Δ1 leu2Δ0 lys2Δ0 ura3Δ0* $ist2^{736\text{-}743\Delta}$ | *This study* |
| MdPY06 | $ist2^{T736A\ T743A}$ | BY4742 | *MATα his3Δ1 leu2Δ0 lys2Δ0 ura3Δ0* $ist2^{T736A\ T743A}$ | *This study* |
| MdPY07 | *cho1Δ* $ist2^{736\text{-}743\Delta}$ | MdPY08 | *MATα his3Δ1 leu2Δ0 lys2ΔΔ0 ura3Δ0 cho1Δ::kanMX* $ist2^{736\text{-}743\Delta}$ | *This study* |

**Table S3.** Plasmids used in this study.

| Plasmid Name | Alias | Description | Origin/Reference |
|---|---|---|---|
| pAK75 | GFP-Ist2 | *IST2$_{pr}$GFP-IST2* (full-length), *CEN, HIS3* (pUG34-Based). | *Kralt et al., 2014* |
| pAK76 | GFP-Ist2RL | *IST2$_{pr}$GFP-IST2* with randomized linker [596-917], CEN, *HIS3* (pUG34-Based). | *Kralt et al., 2014* |
| pAK81 | GFP-IstL240 | *IST2$_{pr}$GFP-IST2* with 100 aa deletion in cytosolic linker, CEN, *HIS3* (pUG34-Based). | *Kralt et al., 2014* |
| pAK77 | GFP-IstL140 | *IST2$_{pr}$GFP-IST2* with 200 aa deletion in cytosolic linker, CEN, His3 (pUG34-Based). | *Kralt et al., 2014* |
| pAK84 | GFP-IstL058 | *IST2$_{pr}$GFP-IST2* with 135 aa deletion in cytosolic, CEN, His3 (pUG34-Based). | *Kralt et al., 2014* |
| pJMD_01 | GFP-Ist2RL$^{[729-747wt]}$ | 57 bp of *IST2* WT [729-747] inserted in the randomized linker of pAK76. | *This study* |
| pJMD_02 | GFP-Ist2[TT->AA] | *T736A T743A* Quikchange mutagenesis of pAK75. | *This study* |
| pJMD_07 | BFP-Ist2 | *IST2$_{pr}$BFP-IST2* full-length, CEN, *HIS3* (pUG34-Based). | *This study* |
| pJMD_08 | BFP-Ist2[705-762] | *IST2$_{pr}$BFP-IST2* [codon705 to 762], CEN, *HIS3* (pUG34-Based). | *This study* |
| pJMD_12 | BFP-Ist2RL$^{[718-751wt]}$ | 102 nt of *IST2* WT [729-747] inserted in the randomized linker of pJMD_21 | *This study* |
| pJMD_21 | BFP-Ist2RL | *IST2$_{pr}$BFP-IST2* with randomized linker [596-917], CEN, HIS3 (pUG34-Based). | *This study* |
| pRS315-Osh6-mCh | pADH1 Osh6-mCherry (L) | *ADH1$_{pr}$OSH6*-mCherry, *CEN, LEU2* (pRS315-based). | *Maeda et al., 2013* |
| pAC100 | pADH1 OSH6-mCherry (U) | *ADH1$_{pr}$OSH*6-mCherry, *CEN, URA3* (pRS315-based); marker swap | *This study* |
| pVA111 | pCYC1 Osh6-mCherry | *CYC1$_{pr}$OSH6*-mCherry, *CEN, URA3* (pRS316-based); *OSH6* subcloned | *This study* |
| pJMD_22 | pADH1$_{pr}$Osh6 *L141A D142A* | Quikchange mutagenesis of pAC100 | *This study* |
| pJMD_23 | pCYC1$_{pr}$Osh6 *L141A D142A* | Quikchange mutagenesis of pVA111 | *This study* |
| pJMD_24 | pADH1$_{pr}$Osh6 *L141K D142A* | Quikchange mutagenesis of pAC100 | *This study* |
| pJMD_25 | pCYC1 Osh6 *L141K D142A* | Quikchange mutagenesis of pVA111 | *This study* |
| pC2$_{Lact}$-GFP | C2$_{lact}$-GFP | GPD$_{pr}$C2$_{Lact}$-GFP, *CEN, URA3* (pRS416-based) | *Yeung et al., 2008* |
| pAC107 | C2$_{lact}$-GFP (LEU) | GPD$_{pr}$C2$_{Lact}$-GFP, *CEN, LEU2* (pRS416-based) | *Lipp et al., 2019* |
| pJMD_26 | crRNA Ist2 736-743 | crRNA targeting *IST2* codons 736-743, *URA*, 2 μ ori, based on pUD628 (Addgene Plasmid #103018) | *This study* |
| pUDC175 | fnCpf1 | *TEF1$_{pr}$FnCPF1, CEN, NatMX* (p414TEF1 Backbone) Addgene Plasmid #103019 | *Swiat et al., 2017* |
| pVA117 | pRS415-ADH$_{pr}$Osh6Δ35-VN | ADH$_{pr}$Osh6Δ35 (Lipp et al., 2019) subcloned into pRS415-VN using SacI/BamHI digestion | *This study* |
| pVA119 | pRS415-ADH$_{pr}$Osh6Δ69-VN | ADH$_{pr}$Osh6Δ69 (Lipp et al., 2019) subcloned into pRS415-VN using SacI/BamHI digestion | *This study* |

**Table S4.** Plasmids constructed for the yeast two-hybrid assay.

| Plasmid Name | Insert | Mutation | Construction |
|---|---|---|---|
| pJMD100 | *OSH4* | - | *OSH4* coding region inserted in the BamHI and XhoI restrictions sites of pGBKT7 (Chien et at., 1991). |
| pJMD101 | *OSH6* | - | *OSH6* coding region inserted in the BamHI and XhoI restrictions sites of pGBKT7 (Chien et at., 1991). |
| pJMD102 | *OSH7* | - | *OSH7* coding region inserted in the BamHI and XhoI restrictions sites of pGBKT7 (Chien et at., 1991). |
| pJMD103 | *OSH6* | *H97D* | Plasmid pJMD103 to pJMD138 were generated by Quikchange site-directed mutagenesis *of pJMD101.* |
| pJMD104 | *OSH6* | *Y135T* | |
| pJMD105 | *OSH6* | *D141K_L142A* | |
| pJMD106 | *OSH6* | *N144D_K145A* | |
| pJMD107 | *OSH6* | *Q146E_Q147A* | |
| pJMD108 | *OSH6* | *Y164F_Y166F* | |
| pJMD109 | *OSH6* | *S171H* | |
| pJMD110 | *OSH6* | *S183A_R184A* | |
| pJMD111 | *OSH6* | *D204N* | |
| pJMD112 | *OSH6* | *K214E* | |
| pJMD113 | *OSH6* | *S244A* | |
| pJMD114 | *OSH6* | *D250N* | |
| pJMD115 | *OSH6* | *E252L* | |
| pJMD116 | *OSH6* | *Y258F* | |
| pJMD117 | *OSH6* | *E267H* | |
| pJMD118 | *OSH6* | *K271E* | |
| pJMD119 | *OSH6* | *Y273K* | |
| pJMD120 | *OSH6* | *S282A* | |
| pJMD121 | *OSH6* | *S282E* | |
| pJMD122 | *OSH6* | *Y290F* | |
| pJMD123 | *OSH6* | *T307A_H308A* | |
| pJMD124 | *OSH6* | *S311A_P312A* | |
| pJMD125 | *OSH6* | *E325A_Y326A* | |
| pJMD126 | *OSH6* | *K333E* | |
| pJMD127 | *OSH6* | *D337K* | |
| pJMD128 | *OSH6* | *H373E* | |
| pJMD129 | *OSH6* | *K375A* | |
| pJMD130 | *OSH6* | *S380A_K381A* | |
| pJMD131 | *OSH6* | *Q403K* | |
| pJMD132 | *OSH6* | *D141K L142A* | |
| pJMD133 | *OSH6* | *D141A L142A* | |
| pJMD134 | *OSH6* | *D141A* | |
| pJMD135 | *OSH6* | *L142A* | |
| pJMD136 | *OSH6* | *D141K L142I* | |
| pJMD137 | *OSH6* | *D141K* | |
| pJMD138 | *OSH6* | *D141A L142D* | |

**Table S4 *continued*.** Plasmids constructed for the yeast two-hybrid assay.

| Name | Insert | Mutation | Construction |
|---|---|---|---|
| pJMD139 | Ist2_1-100 | - | Ist2 coding region from amino acid position 1 to 100 inserted in the BamHI and XhoI restriction sites of pGADT7 (Chien et at., 1991). |
| pJMD140 | Ist2_590-878 | - | Ist2 coding region from amino acid position 590 to 878 inserted in the BamHI and XhoI restriction sites of pGADT7 (Chien et at., 1991). |
| pJMD141 | Ist2_590-703 | - | Ist2 coding region from amino acid position 590 to 703 inserted in the BamHI and XhoI restriction sites of pGADT7 (Chien et at., 1991). |
| pJMD142 | Ist2_762-878 | - | Ist2 coding region from amino acid position 762 to 878 inserted in the BamHI and XhoI restriction sites of pGADT7 (Chien et at., 1991). |
| pJMD143 | Ist2_704-761 | - | Ist2 coding region from amino acid position 704 to 761 inserted in the BamHI and XhoI restriction sites of pGADT7 (Chien et at., 1991). |
| pJMD144 | Ist2_718-751 | - | Ist2 coding region from amino acid position 718 to 751 inserted in the BamHI and XhoI restriction sites of pGADT7 (Chien et at., 1991). |
| pJMD145 | Ist2_729-761 | - | Ist2 coding region from amino acid position 729 to 761 inserted in the BamHI and XhoI restriction sites of pGADT7 (Chien et at., 1991). |
| pJMD146 | Ist2_729-747 | - | Ist2 coding region from amino acid position 729 to 747 inserted in the BamHI and XhoI restriction sites of pGADT7 (Chien et at., 1991). |
| pJMD147 | Ist2_704-727 | - | Ist2 coding region from amino acid position 704 to 747 inserted in the BamHI and XhoI restriction sites of pGADT7 (Chien et at., 1991). |
| pJMD148 | Ist2_704-761 | *S729A* | *Plasmid pJMD148 to pJMD164 were generated by* Quikchange site-directed mutagenesis *of pJMD143.* |
| pJMD149 | Ist2_704-761 | *Y730F* | |
| pJMD150 | Ist2_704-761 | *S729A Y730F* | |
| pJMD151 | Ist2_704-761 | *S729E Y730E* | |
| pJMD152 | Ist2_704-761 | *S729A Y730F T736A* | |
| pJMD153 | Ist2_704-761 | *S729E Y730E T736E* | |
| pJMD154 | Ist2_704-761 | *T743A* | |
| pJMD155 | Ist2_704-761 | *T743E* | |
| pJMD156 | Ist2_704-761 | *Y747A* | |
| pJMD157 | Ist2_704-761 | *Y747E* | |
| pJMD158 | Ist2_704-761 | *T736A T743A* | |
| pJMD159 | Ist2_704-761 | *T736A Y747A* | |
| pJMD160 | Ist2_704-761 | *T736A* | |
| pJMD161 | Ist2_704-761 | *T736E* | |
| pJMD162 | Ist2_704-761 | *T736K* | |
| pJMD163 | Ist2_704-761 | *S744A* | |
| pJMD164 | Ist2_704-761 | *T740A* | |
